# Integrative analyses uncover mechanisms by which aging drives B cell lymphoma

**DOI:** 10.1101/2021.02.23.432500

**Authors:** José P. Castro, Anastasia V. Shindyapina, Alessandro Barbieri, Kejun Ying, Olga S. Strelkova, João A. Paulo, Alexander Tyshkovskiy, Rico Meinl, Csaba Kerepesi, Anna P. Petrashen, Marco Mariotti, Margarita Meer, Yan Hu, Alexander Karamyshev, Grigoriy Losyev, Artur A. Indzhykulian, Steven P. Gygi, John M. Sedivy, John P. Manis, Vadim N. Gladyshev

**Author notes:** These authors contributed equally (José P. Castro, Anastasia V. Shindyapina). ^$^ Current address: i3S, Instituto de Investigação e Inovação em Saúde, Universidade do Porto, 4200-135 Porto, Portugal and Aging and Aneuploidy Laboratory, IBMC, Instituto de Biologia Molecular e Celular, Universidade do Porto, 4200-135 Porto, Portugal.

## Abstract

While cancer is an age-related disease, many cancer studies utilize younger animal models. Here, we uncover how a cancer, B-cell lymphoma, develops as a consequence of a naturally aged system. We show that this malignancy is associated with increased cell size, splenomegaly, and a newly discovered age-associated clonal B-cell (ACBC) population. Driven by exogenous c-Myc activation, hypermethylated promoters and somatic mutations, ACBC cells clonally expand independent of germinal centers (IgM+) and show increased biological age and hypomethylation in partially methylated domains related to mitotic solo-CpGs. Epigenetic changes in transformed mouse B cells are enriched for changes observed in human B-cell lymphomas. Mechanistically, the data suggest that cancerous ACBC cells originate from age-associated B cells, in part involving CD22 protein signaling fostered by the aging microenvironment. Transplantation assays demonstrate that ACBC evolve to become self-sufficient and support malignancy when transferred into young recipients. Inhibition of mTOR or c-Myc in old mice attenuates premalignant changes in B cells during aging and emerges as a therapeutic strategy to delay the onset of age-related lymphoma. Together, we show how aging contributes to B-cell lymphoma through a previously unrecognized mechanism involving cell-intrinsic changes and the aged microenvironment, characterize a model that captures the origin and progression of spontaneous cancer during aging and identify candidate interventions against age-associated lymphoma.

## Introduction

The risk of developing invasive cancer grows exponentially with age starting from midlife ^1^. Although cancer occasionally develops at a young age, the median age of its diagnosis in the US is 66 years old ^2^. Mortality rates for cancer and aging follow the same trend, consistent with the idea that cancer is a disease of aging ^3^. Both normal tissues and cancers accumulate age-related mutations during aging ^4–6^, and age is the main risk factor for cancer development ^7^.

Aging may initiate cancer through a variety of mechanisms, e.g. through genetic or epigenetic alterations and contribution from the microenvironment. One potent oncogene that is also implicated in the rate of aging is *c-Myc* ^8^. Mice overexpressing *c-Myc* develop cancer ^9–11^, whereas *c-Myc* deficient mice are protected from lymphoma and are longer lived ^12^. *c-Myc* also drives cell competition in healthy tissues ^13–15^. Therefore, *c-Myc* may orchestrate age-related clonal expansion, constituting an early step in the transition to cancer. Aberrant DNA methylation is yet another factor thought to initiate age-related cancer as mice deficient in *Tet1/2/3* and *Dnmt3a* develop myeloid leukemias or lymphomas ^16–18^. Likewise, human cancers often exhibit global changes in DNA methylation ^19^. At the same time, many human and mouse tissues accumulate recurrent DNA methylation changes with age, further supporting the idea that aging promotes cancer through epigenetic changes alongside genetic aberrations.

The aging microenvironment is another major driver of cancer transformation ^7^, e.g. acting through chronic inflammation ^20, 21^. Age-associated B cells (ABC) are a prominent constituent contributing a pro-inflammatory environment by overexpressing pro-inflammatory cytokines such as IFNɣ, IL-6, and IL-4 ^22^, in response to TLR stimulation rather than via BCR engagement. ABC accumulate in mammalian aging and disease ^22–27^ and express T-box transcription factor *TBX21* ^24^. However, in contrast to their clear involvement in autoimmunity, infection response and metabolic complications, it remains unclear whether ABC can contribute to age-related clonal selection and cancer.

Most studies uncovering mechanisms of age-related clonal expansion and cancer rely on mouse models derived from known lymphoma targeted gene mutations, rather than wild type mouse models of spontaneous cancer. As a result, they fail to capture the importance of epigenetic and microenvironment shifts for cancer progression associated with aging. Interestingly, C57BL/6 and many other mouse lab strains spontaneously develop and frequently die from B-cell lymphoma ^28, 29^ offering a tool to investigate the full complexity of the origin of a spontaneous age-related cancer. Here, we extensively characterize spontaneous age-related lymphoma in mice, identify a novel IgM+ clonal B cell population named age-associated clonal B cells (ACBC) that transition to cancer without the canonical involvement of germinal centers, offer a mechanism by which ABC as a part of aging microenvironment contribute to cancer transformation and describe therapeutic targets to ameliorate this premalignant ACBC phenotype.

## Results

### B cells increase in size and undergo clonal expansion on the way to age-related lymphoma

We examined age-related changes in the spleen of naturally aged wild type C57BL/6 mice. Old (27 months old) mice were assigned, according to necropsy analyses, to groups free of lymphoma or diagnosed with lymphoma (Fig. 1a). All aged mice, and especially those diagnosed with lymphoma, developed splenomegaly (Fig. 1b, Fig. S1a), which was associated with a prominent increase in the number of B cells and nucleated ones (Fig. 1c, Fig. S1b). Additionally, B cells increased in size, both with age and with the incidence of lymphoma (Fig. 1c), whereas T cells (Fig. S1b) and myeloid cells (Fig. S1c) slightly increased in size.

**Figure 1.**
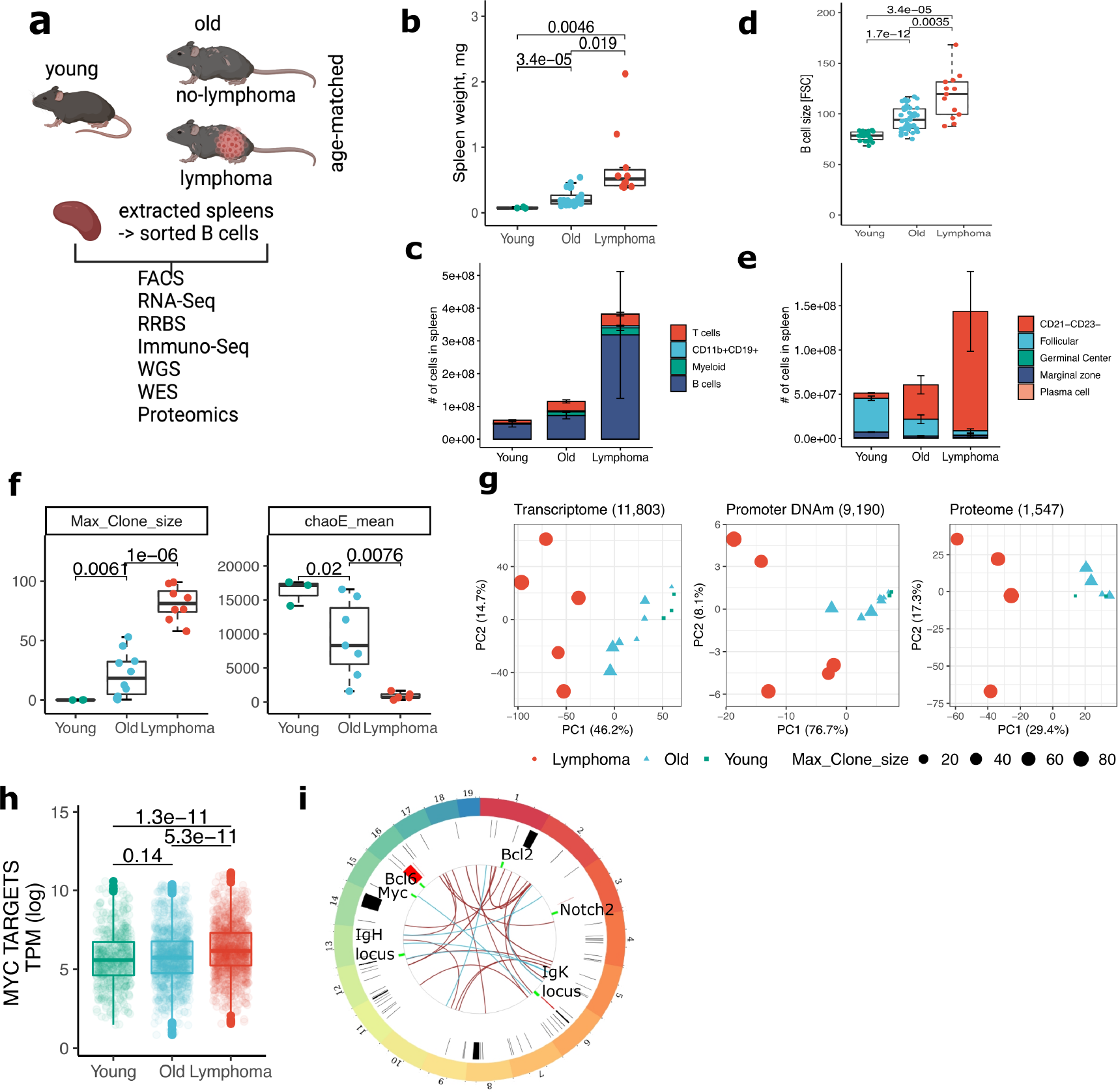
B cells increase in size and undergo clonal expansion driven by somatic mutations and epigenetic alterations during aging. (a) Experimental design for the multi-omics analysis of B cells from young, old and age-matched diseased spleens of C57BL/6 mice. (b) Spleen weights in milligrams collected from the three experimental groups described above. (c) Number of immune cells in the spleens of young, old and old lymphoma-bearing mice as measured by FACS and calculated based on the total number of cells in the tissue. (d) Mean B-cell sizes in young, old and old age-matched lymphoma mice as measured by FACS. Each dot is a mouse. (e) Number of B cells in the spleens of young, old and old lymphoma-bearing mice as measured by FACS and calculated based on the total number of cells in the tissue. (f) Top Ig clone sizes of bulk B cells isolated from young, old and old diseased mice. Clone sizes were estimated by sequencing of CDR3 regions of Ig heavy chains (Max_clone_size, left plot), and CDR3 diversity was estimated with ChaoE metric (ChaoE_mean, right plot). (g) PCA plots of the global transcriptome, promoter DNA methylome and proteome of bulk B cells isolated from young, old and old diseased mice. Numbers indicate the number of features used to produce each plot. (h) Gene expression level of c-Myc targets in bulk B cells from young, old and old diseased mice. (i) Circus plot based on whole genome sequencing of bulk B cells from old and old diseased mice. Red blocks are genomic amplifications, black blocks are genomic deletions, brown and blue lines connect translocation coordinates found for B cells from old and old lymphoma-bearing mice, correspondingly, green blocks depict genes or locus as labeled.

To examine the etiology of enlarged B cells, we analyzed eight common populations of B cells (Fig. S1f). The proportion of follicular (FO) and marginal zone (MZ) B cells significantly decreased with age and most dramatically with lymphoma (Fig. S1d), whereas the proportion and total number of CD21^-^CD23^-^ B cells progressively increased and represented up to 80% of all B cells in the spleens of mice diagnosed with lymphoma (Fig. 1e, Fig. S1d). Of all populations, splenic CD21^-^CD23^-^ B cells showed the most prominent increase in size (Fig. S1e), whereas the size and number of plasma cells and germinal center (Fig. S1e,d) B cells remained unchanged with age and with lymphoma. Thus, the accumulation of CD21^-^CD23^-^ B cells and their increase in cell size largely account for the observed phenotype of enlarged B cells observed in aged mice and mice diagnosed with lymphoma. Importantly, we found that the IgH isotypes in lymphoma cases, mostly dominated by CD21^-^CD23^-^ B cells, were vastly enriched in IgM+ (Fig. S2b, right), which suggests a non-canonical mechanism, excluding germinal center (GC) reaction for age-related lymphoma.

We next tested if enlarged B cells are clonally expanded. We sorted B cells (Fig. S1f) from the spleens of young, old and lymphoma-bearing mice and sequenced variable CDR3 regions of Ig heavy chains to calculate clone sizes (Max_clone_size) and reconstruct Ig diversity (ChaoE_mean). Strikingly, clonal B cells accumulated in substantial numbers with age, and most dramatically in mice diagnosed with lymphoma, where the average clone size was about 80%. Accordingly, significantly lower levels of Ig diversity were observed, suggesting an oligoclonal expansion of B cells with age and cancer (Fig. 1e). Analyses of the spleen and blood data from The Genotype-Tissue Expression (GTEx)^30^ project (Fig. S2l) also revealed a decrease in Ig diversity with age (Fig. S2m,n), as well as a correlation between Ig and genomic clone sizes (Fig. S2o).

Clone size in aged mice positively correlated with B-cell size (R=0.74, p<0.001, Fig. S1g). Accordingly, CDR3 diversity profoundly decreased with age and with the B-cell size increase (Fig. S2a). Top clones did not harbor evidence of immunoglobulin somatic hypermutation within the CDR3 regions in both old and lymphoma cases (Fig. S2b, left), in agreement with IgM positivity excluding GC reactions. This provides evidence for a GC-independent rise of age-related lymphoma in mice. We further sequenced exomes and genomes of B cells isolated from old mice and mice diagnosed with lymphoma and found them to carry hundreds of somatic mutations (Fig. S2f), including highly clonal mutations within cancer-associated genes revealing that age-related B-cell clonal expansion relies on somatic mutations (Supplementary Table 1). Only 5-10% of mutations were shared between regular and enlarged B cells from the same donor, suggesting an independent path for clonal selection of large B cells. B cells isolated from lymphoma-bearing mice on average formed larger clones than their age-matched cancer-free counterparts as measured by variant allele fraction (VAF) (Fig. S2g). Driver somatic mutations were located in the genes commonly mutated in human lymphomas (Supplementary Table 1), indicating partially conserved genetic mechanisms for clonal selection between human and mouse B cells. The top clones determined by CDR3 and exome sequencing were almost identical in size (Fig. S2c), consistent with the idea that the clones carrying driver mutations expand in later stages of B cells maturation cycle (after VDJ rearrangement in the bone marrow) but do not appear to be directly associated with the cytidine deaminase AID off-target alterations suggesting that there are non-GC drivers of age-related lymphoma. B-cell size again significantly correlated with top genomic clone size (Fig. S1g), suggesting that B-cell size is a convenient marker of clonal malignant B cells in mice, especially when surface markers are unknown. Mutational signature deconvolution (Fig. S2h-k) revealed accumulation of SBS29 (73.8% of mutations), SBS42, SBS46, SBS15 and SBS5 signatures in clonal B cells, as well as ID5, DBS7, DBS11, and DBS5, suggesting aging (SBS5, ID5), impaired DNA repair (SBS15, DBS7), APOBEC (DBS11) and possible nucleotide modifications (SBS42, SBS29) as sources of mutations in aging B cells.

To investigate molecular mechanisms driving B cell clonal expansion, we sequenced genomic DNA, mRNA and DNA methylomes, and assessed the proteome of the same set of samples. Principal component analysis (PCA) of genome-wide gene expression, promoter methylation and protein levels (Fig. 1g) revealed that most of the variation between B cell samples is explained by clonal expansion and the experimental group tested (i.e. young vs old vs lymphoma); thus, cancerous B cells exhibit the largest epigenetic perturbation.

Gene targets of c-Myc were overexpressed in lymphoma cases compared to old and young B cells (Fig. 1h). To evaluate whether the phenotype was mediated by *c-Myc* gene translocation or amplification we resorted to whole genome sequencing (WGS) and found no changes in the *c-Myc* genomic region or any other common gene alterations involved in human B cell lymphomas such as *Bcl2* or *Bcl6* (Fig. 1i), supporting non-canonical transformation of B cells.

We further compared genetic and epigenetic alterations of mouse transformed B cells to those found in human lymphomas. We examined gene expression patterns, promoter DNA methylation and somatic mutations in mouse clonal B cells against publicly available datasets of human blood cancer and aging, using breast and pediatric cancer as negative controls and mouse lymphoma signature generated from our dataset as a positive control (Supplementary Table 2). Clonal B cells from aged mice acquired gene expression profiles highly similar to human BCLs, including diffuse large B-cell (DLBCL) and Burkitt lymphoma (BL) (Fig. S1h-left). A deeper analysis revealed that the activation of the c-Myc pathway explained most of the gene expression similarities between human lymphomas and mouse lymphoma (Fig. S1h-right,i). This finding suggests that Myc inflates the transcriptomic similarity between human lymphomas and mouse cancerous B cells. At the same time, promoter DNA methylation of mouse cancerous B cells closely resembled human DLBCL and Follicular lymphoma (FL), but not solid tumors (Fig. S1j). Finally, genes that were commonly mutated in mouse lymphoma were enriched for genes commonly mutated in human DLBCL and FL, but not in solid tumors (Fig. S1k). Overall, the data suggest, at least from this molecular profiling perspective, overlapping epigenetic mechanisms of B-cell clonal selection and malignancy between mice and humans that may account for a more general mechanism of aging driving cancer; thus, aged mice emerge as a viable translational model to study age-related, naturally occurring stochastic processes of B cell malignancy and screen for interventions that preclude B cells from becoming cancerous.

### Age-associated clonal B cells (ACBC) represent a B-cell population with a characteristic premalignant epigenetic program

Our analysis of the epigenomes and application of epigenetic aging clocks revealed that the predicted biological age of clonal B cells ^31^ from lymphoma cases was higher than that of old age-matched B cells even though the mean DNAm was the same (Fig. 2a); the predicted age also correlated with B cell size (Fig. S5b). We further applied mitotic clocks since CGI hypermethylation and heterochromatin hypomethylation are strongly associated with mitotic cell division ^32, 33^. We found substantial hypomethylation in partially methylated domains (PMDs), related to mitotic solo-CpGs ^33^, in aged B cells but most dramatically in transformed B cells (Fig. 2b, left plot) suggesting increased cell divisions. Moreover, using our recently developed mouse multi-tissue transcriptomic clock, we confirmed that B cells isolated from mice diagnosed with lymphoma are predicted to be biologically older compared to young (p=8.3e-5) and to aged B cells (p=0.0049) (Fig. 2b, right plot).

**Figure 2.**
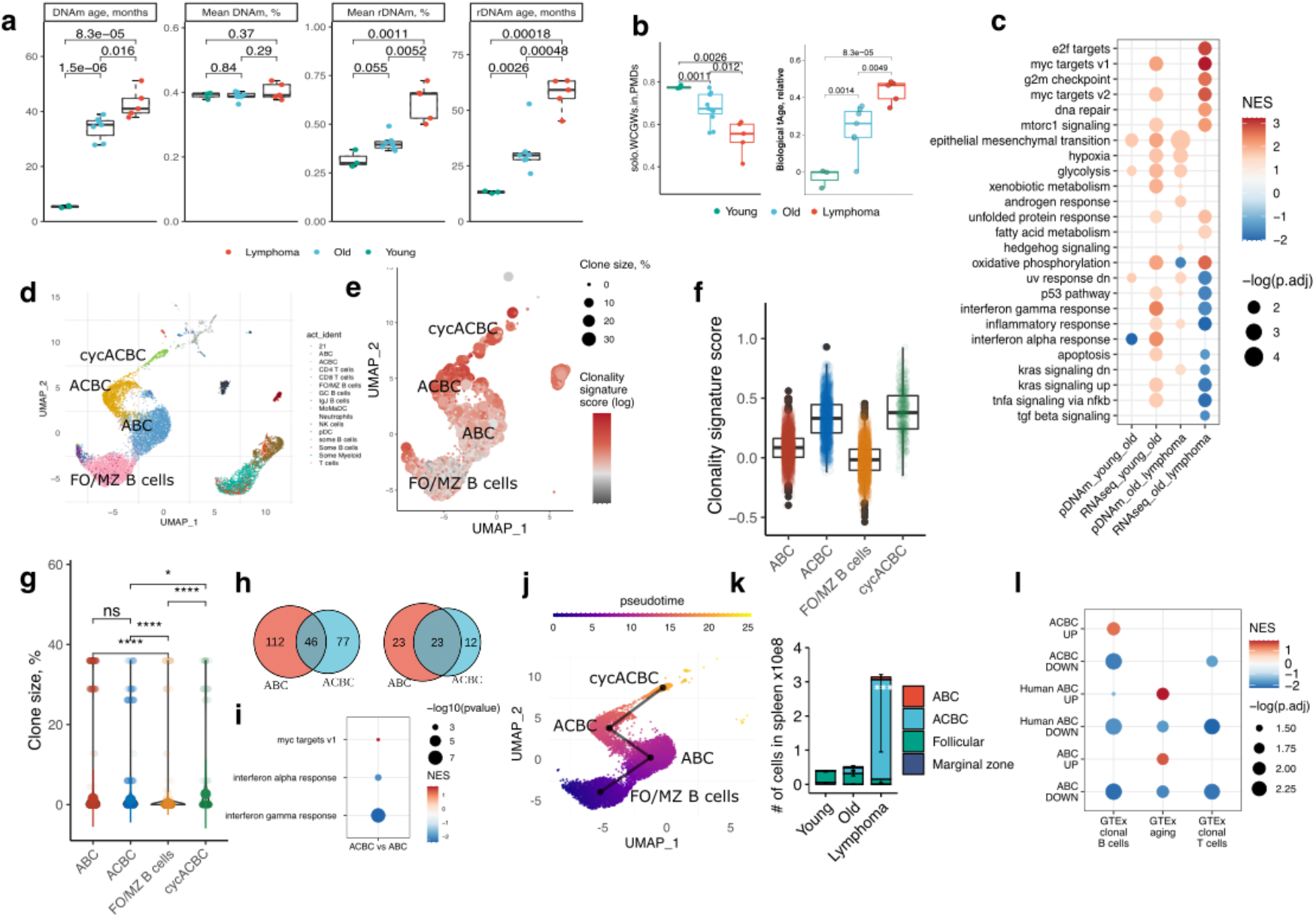
Aged B cells acquire epigenetic programs of malignant cells, and single-cell RNA-seq reveals them as a clonal B-cell population. (a) Mean rDNAm and DNAm (%), rDNAm age and DNAm age (months) of bulk B cells from young, old and lymphoma-bearing mice. (b) Left plot - Mitotic clock for solo-CpGs (WCGW) in partially methylated domains (PMDs) of bulk B cells from young, old and lymphoma-bearing mice. Right plot - Relative biological transcriptomic age (tAge) of young, old and lymphoma-bearing mice. P-values estimated with the independent samples two-tailed t-test are shown in text. (c) MSigDB pathways enriched in differentially methylated promoters in old mice compared to young (pDNAm young old) or in cancerous B cells compared to old (pDNAm old lymphoma), as well as pathways enriched in differentially expressed genes in the same two groups. (d) UMAP plot for scRNA-seq of splenocytes from four old mice combined from two datasets ^36, 37^. (e) UMAP plot from (d) restricted to B cells and colored according to clonality signature score. Dot size is the size of the Ig clone to which this cell belongs. (f) Clonality signature scores for FO/MZ B cells, ABC, ACBC and cycACBC populations from (e), each dot is a cell. (g) Clone sizes estimated for FO/MZ B cells, ABC, ACBC and cycACBC populations from (e), each dot is a cell. P-value was calculated with the Wilcoxon signed-rank test. (h) Overlap between up-regulated and down-regulated genes (FDR<0.05) in ABC and ACBC populations compared to B cells (FO/MZ) in scRNA-seq dataset. (i) MSigDB pathways enriched in genes that are differentially expressed in the ACBC population compared to ABC. (j) Pseudo-time analysis of B cells from the combined publicly available single-cell RNA-seq datasets ^36, 37^. (k) Total number of follicular, marginal zone B cells, ABC and ACBC in the spleens of young, old and old lymphoma-bearing mice. (k) Signatures of mouse ACBC and ABC enriched in differentially expressed genes calculated for blood of GTEx subjects with clonal B cells (GTEx clonal B cells) or T cells (GTEx clonal T cells), as well as for age-related gene expression changes (GTEx aging). **** p-value < 0.0001.

GSEA revealed that cancerous B cells upregulated the expression of *c-Myc* targets*, e2f*, and mTORC1 signaling, central metabolic pathways, and G2/M checkpoint genes (Fig. 2c). While many c-Myc target genes were up-regulated in clonal B cells, expression of many other genes was suppressed, and this pattern was consistent with the patterns of promoter DNA methylation (Fig. 2c, Fig. S5d). In line with what is known about human cancer ^19, 34, 35^, we observed a genome-wide increase in promoter DNA methylation of transformed mouse B cells (Fig. S5d) and also in specific promoters often implicated in lymphoma (Fig. S5f,g).

To investigate the origin of clonal B cells we merged two publicly available single-cell RNA-seq datasets of aged mouse spleens ^36, 37^. Two clusters were specific for aged spleens and were negative for CD21 (*Cr2*) and CD23 (*Fcer2a*) (Fig.2d,S6f) and thus represented CD21^-^CD23^-^ B cells that accumulate with age and with lymphoma diagnosis that we observed with flow cytometry (Fig. 1c). We identified one cluster as previously described Age-associated B cells (ABC), based on the expression of *Tbx21* and CD11c (*Itgax*)^24^ (Fig. 2d) and application of gene expression signatures of ABC cells from publicly available mouse datasets (Fig. S6a). This population has been found in the spleen of human subjects with DLBCL ^38^To test which B cell population was clonal, we estimated the clonality signature score for each cell based on a correlative score we extracted from bulk RNA-seq by correlating gene expression with the largest clone size of B-cell sample (Supplementary Table 3). The other cluster of age-related B cells had the highest scores among aged B cells (Fig. 2e). Reconstruction of CDR3 regions of Ig *kappa* chains in individual cells revealed that clonal B cells belonged to one of the age-emergent B-cell clusters (Fig. 2f,g). We named these cells as age-associated clonal B cells (ACBC). GSEA analysis of the ACBC cluster revealed a significant up-regulation of c-Myc targets compared to ABC cells and a decrease in IFN pathways, suggesting that ACBC cells might become insensitive to immune responses (Fig. 2h). Notably, the clonality signature score significantly correlated with Igk clone size in both datasets (Fig. S6e), and thus it might be relevant to identify clonal B cells in other single-cell spleen datasets.

We observed that both age-related clusters of B cells, ABC and ACBC, shared many deregulated genes compared to the rest of splenic B cells (FO/MZ) (Fig. 2i). Moreover, ACBC and ABC shared CDR3 Igk sequences predicted from single-cell RNA-seq (Supplementary Table 4). Because ABC are present alongside the ACBC population in every flow cytometry data and in scRNA-seq datasets, but not the other way around, ACBC appear to be a successor cell type from ABC, providing a mechanistic link between aging and cancer. Gene expression based-pseudotime analysis also predicted the ACBC to originate from ABC which in turn had their origin in FO B cells (Fig. 2j). Another population that appears with age and seems to originate from ACBC was enriched in cell cycle mitotic genes and is likely to represent cyclic ACBC (cycACBC) (Fig. S6b,c).

To differentiate and validate our findings with flow cytometry, we predicted CD29 as a positive marker for ABC and ACBC populations, and CD24a as a specific positive marker for ACBC using differential expression analysis (Fig. 4g). qRT-PCR data confirmed that our antibody panel differentiated ABC from ACBC, as FACS-sorted ABC cells (gated as CD19^+^CD21^-^CD23^-^CD29^+^CD24^-^) overexpressed *Tbx21*, and ACBC (gated as CD19^+^CD21^-^CD23^-^CD29^+^CD24^+^) overexpressed *c-Myc* (Fig. S6f). Using this new antibody panel (Fig. S6f), we further revealed that ACBC accumulated with age in the mouse spleen and blood and to a larger extent in age-related lymphoma cases (Fig. 2k). Based on forward scatter, ACBC were larger than follicular B cells (Fig. S6g,h).

Using scRNA-seq datasets we established an ACBC signature and found mouse peripheral tissues to be strongly enriched in it with age suggesting that ACBC infiltrate a multitude of peripheral tissues including fat, liver, lung and kidney (Fig. S4). The same signature was enriched in blood of healthy human subjects from the GTEx dataset (Fig. 2i), suggesting that ACBC also accumulate in humans. Indeed, elderly subjects had increased clonal B cells that are enriched in ABC and ACBC signatures (FIg. S3a,b) as demonstrated by our analysis of publicly available dataset ^39^ of sc-RNA-seq data coupled with VDJ-seq. Consistent with clonal mouse B cells, 31% of predicted human ACBC cells were IgM+ (Fig. S3c).

Based on single-cell and bulk RNA-seq, we conclude that clonal B cells in old mice diagnosed with lymphoma are predominantly represented by an ACBC population, that ABC likely to give rise to ACBC, and that c-Myc is a potential driver of B-cell clonal expansion and cell size increase with age. We further established a panel of markers to differentiate clonal ACBC from ABC using flow cytometry.

### Clonal B cells (ACBC) are the predictors of lifespan, originate with the support of an aging microenvironment and become self-sufficient over time

We next investigated interaction among B cells in the aged spleen to shed light on the origin of ACBC and the cell size increase. CellChat predicted ABC to interact with themselves and with ACBC (Fig. 3a) through a few receptor-receptor interactions, including the one through CD22 as shown in their incoming and outgoing signaling pathways (Fig. 3b). This prediction raised the possibility of ABC-ABC interaction being in the origin of ACBC formation. We further found strong ABC interaction with CD8+ and CD4+ T cells (Fig. S6d), which may be a sign of an extra-follicular T cell dependent mechanism of ABC differentiation, as shown previously ^38^.

**Figure 3.**
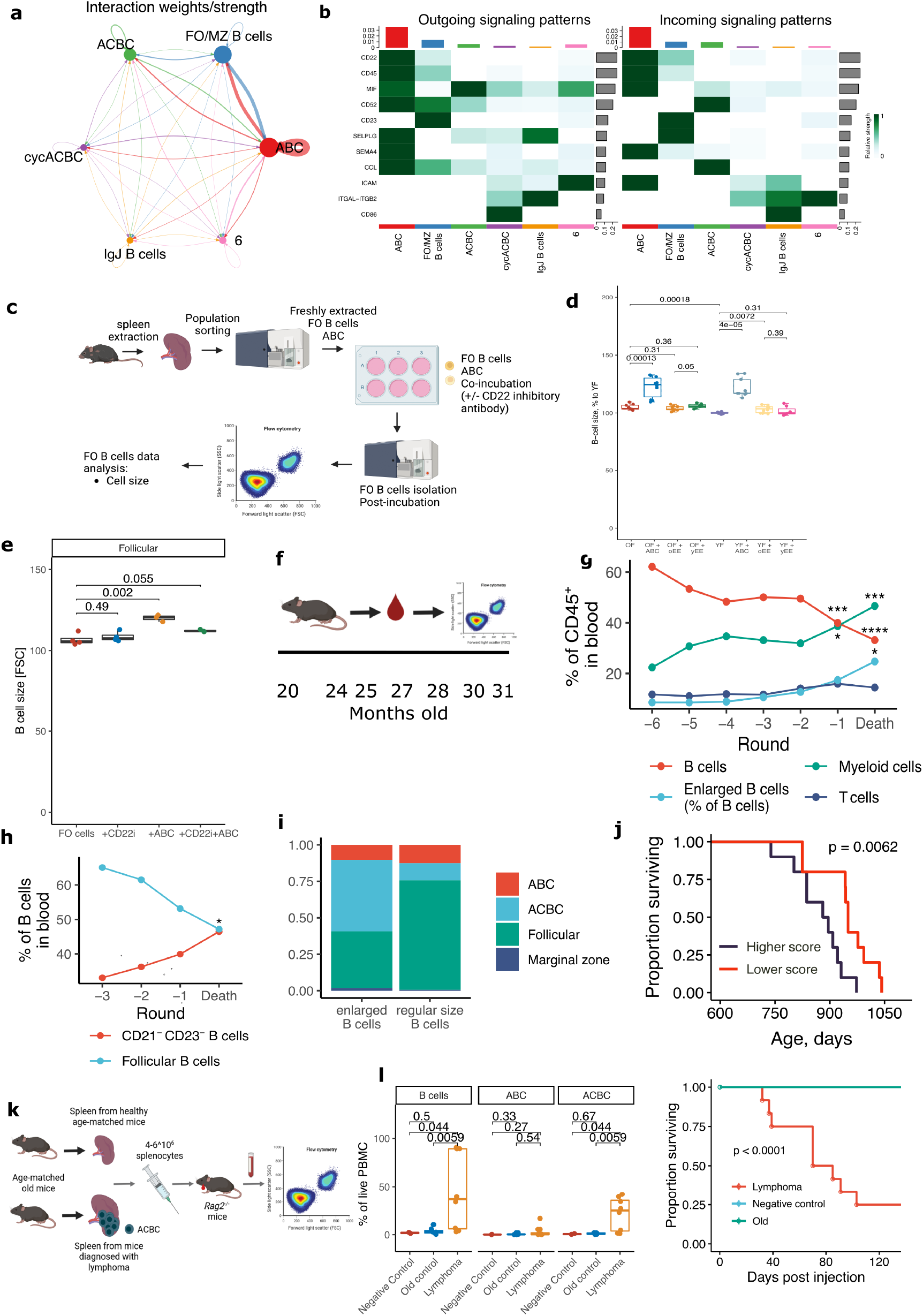
Longitudinal analysis and transplantation experiments reveal ACBC cells as a cancerous B-cell population that leads to animal fatalities, and co-culture experiments point to their origin. (a) Predicted strength of molecular interactions between B-cell populations from combined scRNA-seq dataset used in Fig. 2. (b) Heatmaps of predicted incoming and outgoing signals between various B cell populations in aged mouse spleens. (c) Scheme for co-culture and CD22 inhibition experimental layout. (d) Mean cell size measured by FACS in young (YF) and old follicular (OF) B cells incubated with i) ABC from old spleens, ii) non-B cells immune cells (CD45^+^CD19^-^) from young spleens (yEE), or iii) non-B cells immune cells (CD45^+^CD19^-^) from old spleens (oEE). Results shown are from three independent experiments, 3-4 technical replicas per experiment. (e) Mean size of B cells cultured i) alone, ii) with ABC, iii) with CD22 inhibitory (CD22i) monoclonal antibody or iv) with ABC and with CD22 inhibitory monoclonal antibody. (f) Longitudinal blood collection from tail veins of 20 C57BL/6 female mice across seven timepoints. (g) Changes in cellular composition of blood from old mice aligned to the last blood collection before animal death (e.g., Death is the last blood collection before mice death, -1 is the time point before that last blood collection). (h) Same as in (g) but for follicular and CD21-CD23-B cells. (j) B-cell populations present in regular-sized B cells or enlarged B cells in old mouse spleens gated based on forward scatter. (j) Survival curve of mice with higher and lower presence of CD21^-^CD23^-^ and enlarged B cells in 27-month-old mice. (k) Scheme for the transplantation experiments for splenocytes from old mice diagnosed with lymphoma or healthy age-matched old mice (Old control) into young *Rag2^-^*^/-^ mice. (l) Percentage of B-cell populations to live CD45^+^ cells measured in blood of *Rag2^-^*^/-^ mice one month post i.p. injection of splenocytes or left without injections (negative control) (left plot). (m) Survival of *Rag2^-^*^/-^ mice after transplantation of splenocytes from mice diagnosed with lymphoma or old healthy mice, or mice injected with saline. P-values for survival curves were calculated with a log-rank test (right plot).

To confirm these observations, we performed co-culture experiments exposing FO cells to ABC cells and CD45+CD19-splenic cells (Fig. 3c). Follicular B cells increase in size the presence of ABC *in vitro* (Fig. 3d) and overexpress *Myc* (Fig. S7d). To validate CellChat (Fig. 3b) predictions, we further co-cultured FO with ABC cells in presence of monoclonal CD22 antibody. Interestingly, FO cells increase in size less when co-incubated with ABC and a CD22 inhibitory antibody compared to co-culture with ABC alone (Fig. 3e). This finding suggests that ABC trigger cell size increase which involves c-Myc activation and CD22 signaling.

Next, in a longitudinal study, we assessed whether an earlier accumulation of clonal B cells was associated with a shorter lifespan in mice (Fig. 3f). Longitudinal blood analysis (Fig. 3g) established that enlarged B cells accumulate in the mouse blood starting at the age of 28 months and that myeloid bias started at about 25 months of age (Fig. S7b,c). The trajectory of blood age-related changes became more consistent when the mice were aligned by the death rather than by the age which demonstrates that B-cell size increase is more predictive of lifespan than of chronological age (Fig. 3h). Since this analysis was done before the ACBC flow panel was established, we used percentage of large B cells as a proxy for proportion of clonal B cells, and later confirmed that enlarged B cells are predominantly composed of ABC in old animals and of ACBC in mice diagnosed with lymphoma (Fig. 3i).

To test whether ACBC can survive outside of the aging environment we injected spleen extracts from mice diagnosed with lymphoma (mostly consisting of ACBC) into immunodeficient *Rag2*^-/-^ mice (Fig. 3k). One month after injection, we detected significant presence of ACBC in the blood of most recipients, that received splenocytes from lymphoma-bearing mice but not age-matched controls (Fig. 3l, left). Nine out of 10 mice injected with splenic extracts (mostly containing ACBC) from mice diagnosed with lymphoma died within 103 days, while none of the control mice died (Fig. 3l, right) validating the causal relationship between ACBC and premature death observed in longitudinal data and malignancy of ACBCs.

Altogether, our data suggest that aged follicular B cells enlarge in size to become ABC through CD22 signaling, lead to *c-Myc* expression harboring advantageous somatic mutations and thus selecting out cells with the c-Myc overexpression transcriptional program in transitioning from ABC into ACBC. We thus hypothesized that the aging microenvironment, especially ABC cells, apply selective pressure on aged B cells by activating their c-Myc, and selecting out genetic and epigenetic states that favor c-Myc overexpression. That process might give rise to clonal *Myc*-overexpressing B cells, the ACBC (Fig. 3i) ^40, 41^. Over time, ACBC cells evolve into self-sufficient malignant cells manifested in down-regulation of inflammatory pathways, apoptotic and p53 pathways, and up-regulation of cell division mitotic genes (Fig. 2c; Fig. S6b,c) that thrive outside of the aging microenvironment as evident from transplantation study.

### c-Myc deficiency and rapamycin attenuate age-related markers and clonal expansion of B cells

To test a causal role of c-Myc in age-related clonal expansion, we analyzed c-Myc heterozygous mice (Myc+/-, Fig. 4a) ^12^ and mice overexpressing c-Myc specifically in B cells (Eμ-Myc) ^10^. We found that c-Myc haploinsufficiency (Fig. S8a) attenuated splenomegaly (Fig. S8b), the increase in total B-cell size, the ACBC size (Fig. 4b,c), and the accumulation of ACBC (Fig. 4d). Moreover, Myc+/- B cells preserved a more diverse Ig repertoire with age (Fig. 4e). Other immune cell types were not affected by c-Myc deficiency (Fig. S8d,j) suggesting a cell-type and stage specific role of c-Myc in B-cell aging and malignancy.

**Figure 4.**
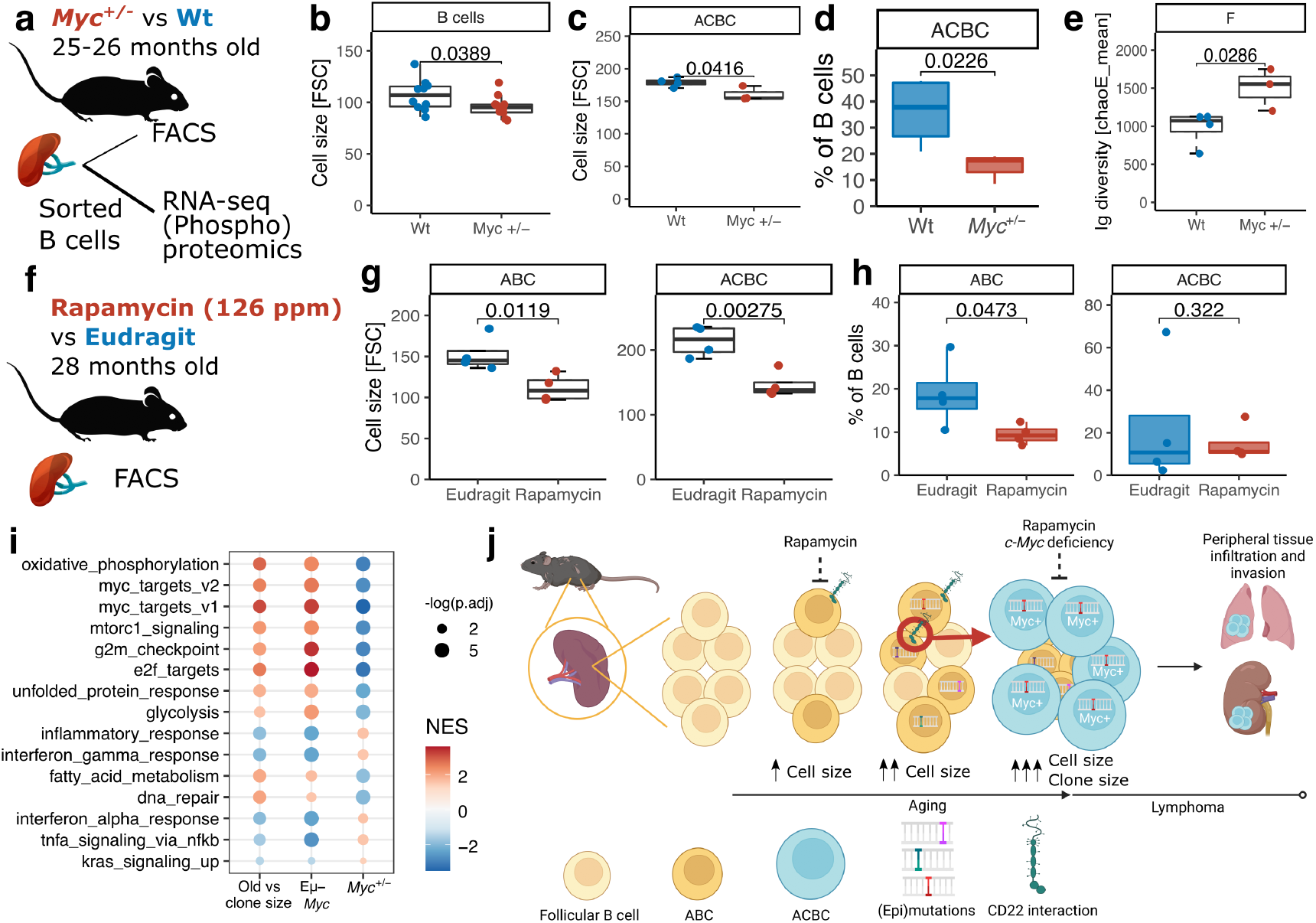
Myc deficiency and rapamycin treatment inhibit cell size increase and age-related clonal expansion of B cells. (a) Analysis of wild type (Wt) and c-Myc haploinsufficient (*Myc*^+/-^) siblings. (b) Mean size of B cells isolated from the spleens of old Wt and *Myc*^+/-^ mice. (c) Mean size of ACBC isolated from the spleens of 26-month-old Wt and *Myc*^+/-^ mice. (d) Percentages of ACBC to total B cells (CD19^+^) in 26-month-old Wt and *Myc*^+/-^ mice. (e) Ig diversity (chaoE_mean) of splenic B cells from 26-month-old Wt and *Myc*^+/-^ mice. (f) Analysis of 27-month-old C57BL/6 mice treated with 126 ppm of encapsulated rapamycin or encapsulating material for 4 months. (g) Mean cell size of ACBC from spleens of old mice treated with rapamycin or control diets. (h) Percentage of ABC and ACBC to total splenic B cells (CD19^+^) of old mice treated with rapamycin or control diets. (i) MSigDB pathways enriched in differentially expressed genes: 1) in splenic B cells with clonal expansion compared to non-clonal B cells in old C57BL/6 mice, 2) in splenic B cells from young Eμ-*Myc* mice compared to wild type controls, and 3) in B cells from old *Myc*^+/-^ mice compared to their wild type siblings. P-values were calculated with a one-sided Student t-test. (j) Proposed mechanism for the development of ACBC and age-related B-cell lymphoma.

At the same time, splenic B cells from Eμ-*Myc* mice (Fig. S8k) were larger in the spleen (Fig. S8l) and bone marrow (Fig. S8m) and exhibited reduced diversity of the Ig repertoire than age-matched non-carriers (Fig. S7n). Together, these data strongly suggest that c-Myc regulates cell size and fosters clonal expansion of B cells. We further subjected 25-old B6 mice to rapamycin, which inhibits mTOR and c-Myc transcriptional activity ^42^, and found that it attenuated the cell size increase seen in ABC and ACBC (Fig. 4g) and the accumulation of ABC (Fig. 4h) following 3 months of treatment.

Bulk RNA-seq of sorted B cells from both genetic mouse models demonstrated that clonal B cells from aged mice and Eμ-*Myc* B cells exhibit similar expression pattern alterations, whereas *c-Myc*-deficient B cells were opposite to both (Fig. 4i). Genes that belong to *c-Myc* targets, metabolic and inflammation pathways cooperatively respond to changes in *c-Myc* level, and thus the majority of changes in the transcriptome of clonal B cells that accumulate with age can be attributed to *c-Myc* overexpression. These data point to therapeutic targets in preventing the transformation of aged B cells towards malignancy.

## Discussion

To understand how cancer develops in the context of aging we studied a commonly used mouse strain, C57BL/6 mice, which spontaneously develops B-cell lymphoma with age ^28, 29^. We found that mouse clonal B cells acquire genomic, gene expression and DNA methylation patterns that are enriched in human lymphomas, in particular, follicular and diffuse-large B-cell lymphomas, but not in solid tumors. This unexpected result points to evolutionary conserved mechanisms of age-related B-cell malignancy at the genetic and epigenetic levels. Further studies may investigate if other spontaneous cancers observed in mice evolve similarly to human cancers with age.

We further uncovered a population of age-associated clonal B cells, the ACBC, that accumulate with age and evolve into cancerous B cells over time. c-Myc emerged as a major driver of B-cell clonal expansion with age as confirmed by transcriptome studies and mouse models. Our DNA methylation analyses revealed hypermethylated promoters in clonal B cells that silenced gene expression similarly to those in human cancer ^19, 34, 35^. One of the silenced genes was *Id3,* which is a human and mouse tumor suppressor ^43–46^, providing an example of how age-related changes in DNA methylation can contribute to B-cell malignancy. At the same time, ACBC lack somatic hypermutations in CDR3 regions and are IgM+, and thus do not seem to undergo Ig selection in germinal centers following the canonical pathway for lymphoma transformation, and although the mouse and human molecular landscapes are similar, mouse lymphoma does not seem to require the canonical SHM reaction in the germinal center and *c-myc* translocation but rather to be dependent on a set of previously unrecognized events such as ABC-ABC interaction and CD22 signaling. Together, these results suggest that the age-related DNA methylome, somatic mutations and activated c-Myc provide selective advantage to B cell clones in mice rather than the canonical Ig-selection in humans.

ABC shared the majority of deregulated genes and Ig clones with ACBC, suggesting that age-associated B cells give rise to clonal B cells in mice. In humans, ABC accumulate in elderly patients that are at higher risk of B-cell lymphomas, such as in human infections ^47, 48^, and certain autoimmune diseases ^25, 27, 49^ susceptible to diffuse-large B-cell lymphoma ^50^. We further found a population of clonal B cells enriched for ACBC signature in humans over 50 years of age. Thus, our findings of ABC giving rise to ACBC in mice are relevant in the context of aging and cancer in humans.

Although activation of *c-Myc* is a consequence of genetic perturbation in many cancers ^51^, there were no genetic alterations that can explain up-regulation of *c-Myc* in mouse B cells during aging. Instead, exposure of follicular B cells to ABC was sufficient to elevate c-Myc expression through CD22 signaling. Despite ACBC arising from the aging microenvironment that contains ABC, malignant ACBC are transplantable into young immunodeficient mice, demonstrating that surrounding signals are essential to initiate rather than to support cancer progression.

Finally, we demonstrated that two interventions, a genetic (c-Myc deficiency) and a pharmacological (rapamycin) that potently extend mouse lifespan ^12, 52^ attenuate malignant phenotypes of B cells. Our longitudinal study further revealed that the accumulation of clonal B cells stratifies aged mice into long- and short-lived phenotypes. It is possible that c-Myc deficiency and rapamycin (and possibly other longevity interventions) extend mouse lifespan, at least in part, by delaying age-related lymphoma. Furthermore, based on our data of conserved mechanisms of B-cell malignancy between human and mouse, mouse longevity interventions emerge as therapeutic strategies to delay B-cell malignancy in elderly patients.

Overall, our work demonstrates the relevance of epigenetic and genetic mechanisms behind spontaneous age-related mouse lymphomas to human lymphomas and provides mechanistic data that aged mice is a convenient model to tease out the role of microenvironment, epigenetics and genetics in age-emergent cancer. Moreover, we provide evidence that age-related mechanisms that drive lymphoma originate not from the canonical AID and germinal center reactions but from a compilation of human-like epigenetic and genetic selection and signals of the aging microenvironment, including the accumulation of ABC.

## Experimental Procedures

### Animals

C57BL6/JNia mice were obtained from the National Institute on Aging Aged Rodent Colony. Only female mice were used in the present study. 6-wk-old female Eμ-*Myc* mice and control non-carriers were purchased from The Jackson Laboratory (cat. #002728). Animals were euthanized with CO_2_. Spleens were harvested and stored in cold PBS until analysis, and liver samples were immediately frozen in liquid nitrogen and stored at - 80°C. For bone marrow analyses, bones were stripped of the muscle in cold PBS, cut from both ends, and cells were aspirated with 5 ml of cold PBS. The aspirates were filtered through 40 µm Falcon Cell Strainers (Corning). Spleens from 24-27 months old Myc haploinsufficient (*Myc* ^+/-^) mice and their wild type siblings were collected at Brown University and transferred on cold PBS to Harvard Medical School for further analysis. Mice were subjected to encapsulated rapamycin (Rapamycin Holdings; 126 ppm of active compound) or to encapsulating material of the same concentration (Rapamycin Holdings) in 5053 diet (TestDiet) and were fed *ad libitum*. For transplantation experiments, total splenocytes were cryopreserved in 10% DMSO FBS. Recovered splenocytes were counted and 2-6 million cells in 100 ul sterile saline solution (Sigma) were i.p. injected into 2-3 months old *Rag2*^-/-^ mice. *Rag2*^-/-^ mice were housed in the Manis laboratory. All experiments using mice were performed in accordance with institutional guidelines for the use of laboratory animals and were approved by the Brigham and Women’s Hospital and Harvard Medical School Institutional Animal Care and Use Committees.

### Flow cytometry and sorting

mAbs used for staining included: anti-CD19 [6D5], anti-CD43 [S11], anti-IgM [RMM-1], anti-CD11b [M1/70], anti-B220, anti-CD80 [16-10A1], anti-CD3 [17A2], anti-Fas [SA367H8], anti-CD138 [281-2], anti-CD23 [B3B4], anti-CD21 [7E9], anti-CD24 [M1/69], anti-CD29 [HMβ1-1], anti-CD93, and anti-CD45 [30-F11] (all from Biolegend) with fluorophores as in Supplementary table 7. Dead cells were excluded by DAPI staining. Data was collected on a Cytek DXP11 and analyzed by FlowJo software (BD). Cells were sorted on BD Aria Fusion. Spleens were gently pressed between microscopy slides to get single-cell suspensions. One ml of cell suspensions were 1) incubated with 14 ml of red blood lysis buffer for 10 min on ice, 2) centrifuged at 4°C, 250 g for 10 min, 3) washed once with 1 ml of FACS buffer (1% FBS in PBS), 4) stained in 100 ul of AB solution (2 ng/ul of each AB) at 4°C for 20 min protected from light, 5) washed again, 6) filtered into tubes with cell strainer snap cap (Corning), and 7) analyzed with flow cytometry or FACS-sorted into 1 ml of FACS buffer. Surface markers of ABC and ACBC were selected based on: 1) differential up-regulation in both populations in scRNA-seq, 2) annotation as surface protein in GO, 3) differentially up-regulated in clonal B cells in bulk RNA-seq data (Supplementary table 3). It yielded a few candidates, of which we chose CD29 (*Itgb1*) as a positive marker for both age-related populations, and CD24 to distinguish ACBC from ABC.

### RNA sequencing and qRT-PCR

Total RNA, DNA and proteins were extracted from fresh or snap-frozen FACS-sorted 2-5 million splenic B cells using AllPrep DNA/RNA/Protein Mini Kit (Qiagen) following the manufacturer’s instructions. RNA was eluted with 42 ul of RNAse-free water. RNA concentration was measured with Qubit using the RNA HS Assay kit. Libraries were prepared with TruSeq Stranded mRNA LT Sample Prep Kit according to TruSeq Stranded mRNA Sample Preparation Guide, Part # 15031047 Rev. E. Libraries had been quantified using the Bioanalyzer (Agilent), and were sequenced with Illumina NovaSeq6000 S4 (2x150bp) (reads trimmed to 2x100bp) to get 20M read depth coverage per sample. The BCL (base calls) binary were converted into FASTQ using the Illumina package bcl2fastq. Fastq files were mapped to mm10 (GRCm38.p6) mouse genome. and gene counts were obtained with STAR v2.7.2b ^53^. Myc targets for Figure 2c were taken from the hallmark set of genes ‘MYC_TARGETS_V1’ from msigdb database. For quantitative RT-PCR, RNA samples were normalized by DNA concentration isolated from the same sample, then 2 ul of normalized RNA were mixed with iTaq Universal SYBR (Bio-Rad) and primers for *c-Myc* (F: TTCCTTTGGGCGTTGGAAAC, R: GCTGTACGGAGTCGTAGTCG), *Actb* (F: GGCTGTATTCCCCTCCATCG, R: CCAGTTGGTAACAATGCCATGT) or *Gapdh* (F: AGGTCGGTGTGAACGGATTTG, R: TGTAGACCATGTAGTTGAGGTCA) for the final volume of 10 ul, and loaded into Multiplate 96-Well PCR Plates (Bio-rad). Data was acquired for 40 cycles on Bio-Rad C1000, CFX96 Thermal Cycler. Each sample was loaded in duplicate.

### Whole exome sequencing

Total DNA was extracted with AllPrep DNA/RNA/Protein Mini Kit (Qiagen) following the manufacturer’s instructions. DNA was eluted with 100 ul of EB. DNA concentration was measured with Qubit using DNA BR Assay kit. Libraries were prepared with Agilent SureSelect XT Mouse All Exon according to SureSelectXT Target Enrichment System for Illumina Version B.2, April 2015. Libraries had been quantified using the Bioanalyzer (Agilent), and were sequenced with Illumina NovaSeq6000 S4 (2x150bp) (reads trimmed to 2x100bp) to get 100X throughput depth (roughly 50X on-target) coverage per sample. The BCL (base calls) binary were converted into FASTQ using the Illumina package bcl2fastq. Genomic reads were mapped to the GRCm38.p2 mouse genome assembly using BWA-MEM v0.7.15-r1140 ^54^ and sorted using Samtools v1.6 ^55^. Somatic mutations were called with Mutect2 from the GATK package v4.1.8.0 using liver as matched control; high quality variants were selected with GATK FilterMutectCalls ^56^. Variants were annotated with the Ensembl Variant Effect Predictor v100.2 (with option --everything) ^57^. Only the variants that 1) were supported by 5 reads or more (DP.ALT>4), and 2) were in positions covered by 30 reads or more (DP.ALT+DP.REF>29), were taken for VAF analysis. Signatures of somatic mutations were extracted and deconvoluted using Sigproextractor v.1.0.20 with 100 nmf replicates.

### Whole genome sequencing

DNA was prepared and sequenced as described in the whole exome sequencing section.

### Proteomics

Tandem mass tag (TMTpro) isobaric reagents were from ThermoFisher Scientific (Waltham, MA). Trypsin was purchased from Pierce Biotechnology (Rockford, IL) and LysC from Wako Chemicals (Richmond, VA). Samples were prepared as described previously ^58, 59^. Briefly, cell pellets were syringe-lysed in 8 M urea complemented with protease and phosphatase inhibitors. Samples were reduced using 5 mM TCEP for 30 min and alkylated with 10 mM iodoacetamide for 30 min. The excess of iodoacetamide was quenched with 10 mM DTT for 15 min. Protein was quantified using the BCA protein assay. Approximately 50 µg of protein were chloroform-methanol precipitated and reconstituted in 100 µL of 200 mM EPPS (pH 8.5). Protein was digested using Lys-C overnight at room temperature followed by trypsin for 6h at 37°C, both at a 100:1 protein:protease ratio. After digestion, the samples were labeled using the TMTpro16 reagents for 90 min, the reactions were quenched using hydroxylamine (final concentration of 0.3% v/v). The samples were combined equally and subsequently desalted. We enriched phosphopeptides from the pooled TMT-labeled mixtures using the Pierce High-Select Fe-NTA Phosphopeptide Enrichment kit (“mini-phos”) ^59, 60^ following manufacturer’s instructions. The unbound fraction was retained and fractionated using basic pH reversed-phase (BPRP) HPLC. Ninety-six fractions were collected and then consolidated into 12 which were analyzed by LC-MS3 ^61^.

All data were collected on an Orbitrap Fusion Lumos mass spectrometer coupled to a Proxeon NanoLC-1000 UHPLC. The peptides were separated using a 100 μm capillary column packed with ≈35 cm of Accucore 150 resin (2.6 μm, 150 Å; ThermoFisher Scientific). The mobile phase was 5% acetonitrile, 0.125% formic acid (A) and 95% acetonitrile, 0.125% formic acid (B). For BPRP fractions, the data were collected using a DDA-SPS-MS3 method with online real-time database searching (RTS)^62^ to reduce ion interference ^63, 64^. Each fraction was eluted using a 90 min method over a gradient from 6% to 30% B. Peptides were ionized with a spray voltage of 2,600 kV. The instrument method included Orbitrap MS1 scans (resolution of 120,000; mass range 400−1400 m/z; automatic gain control (AGC) target 2x105, max injection time of 50 ms and ion trap MS2 scans (CID collision energy of 35%; AGC target 1x104; rapid scan mode; max injection time of 120 ms). RTS was enabled and quantitative SPS-MS3 scans (resolution of 50,000; AGC target 2.5x105; max injection time of 250 ms). Raw files were first converted to mzXML. Database searching included all mouse entries from UniProt (downloaded March 2020). The database was concatenated with one composed of all protein sequences in the reversed order. Sequences of common contaminant proteins were also included. Searches were performed using a 50ppm precursor ion tolerance and 0.9 Da (low-resolution MS2) or 0.03 Da (high-resolution MS2) product ion tolerance. TMTpro on lysine residues and peptide N termini (+304.2071 Da) and carbamidomethylation of cysteine residues (+57.0215 Da) were set as static modifications, and oxidation of methionine residues (+15.9949 Da) was set as a variable modification. For phosphopeptide analysis, +79.9663 Da was set as a variable modification on serine, threonine, and tyrosine residues.

PSMs (peptide spectrum matches) were adjusted to a 1% false discovery rate (FDR) ^65, 66^. PSM filtering was performed using linear discriminant analysis (LDA) as described previously^68^, while considering the following parameters: XCorr, ΔCn, missed cleavages, peptide length, charge state, and precursor mass accuracy. Protein-level FDR was subsequently estimated. Phosphorylation site localization was determined using the AScore algorithm ^67^. A threshold of 13 corresponded to 95% confidence that a given phosphorylation site was localized.

For reporter ion quantification, a 0.003 Da window around the theoretical m/z of each reporter ion was scanned, and the most intense m/z was used. Peptides were filtered to include only those with a summed signal-to-noise ratio ≥100 across all channels. For each protein, the filtered signal-to-noise values were summed to generate protein quantification values. To control for different total protein loading within an experiment, the summed protein quantities of each channel were adjusted to be equal in the experiment. For each protein in a TMTpro experiment, the signal-to-noise was scaled to sum to 100 to facilitate comparisons across experiments.

Spectral counts values were analyzed with R in Rstudio. Proteome and phosphoproteome data was normalized using the RLE method and log transformed using the *edgeR* package ^68^. Values for phospho sites were normalized to corresponding protein level and differentially changed sites were calculated with the limma package. Ranked phospho sites were then assessed for enrichment for targets of mouse kinases using PTM-SEA resources ^69^ and kinact software ^70^.

### Single-cell RNA sequencing analysis

To identify genes differentially expressed in this newly identified B cell cluster and other B cells, we downloaded the single cell RNA seq data from Calico’s murine aging cell atlas (https://mca.research.calicolabs.com/data, spleen single-cell count data, filtered) ^36^. Preprocessing of the downloaded data was performed using scanpy ^71^. We first removed cells with a high (>0.05) percentage of mitochondrial reads. Cells not annotated as B cells were also removed. We then normalized the read counts by total reads number per cell and multiplied by a rescaling factor of 10000. Normalized reads were log transformed after adding a pseudo-count of 1. We scaled the log-transformed data to unit variance and zero mean and clipped maximum value to 10. After the above data preprocessing, we selected the cells corresponding to the age-related B-cell cluster, which was C130026I21Rik^+^Apoe^+^Cr2^-^Fcer2a^-^. For each cell, we used a linear combination of the RNA level of these four marker genes to calculate a score (score = C130026I21Rik + Apoe - Cr2 - Fcer2a), and within this cluster we selected the ABC cluster that is Tbx21^+^ and another one that is Myc^+^ using the same linear system. We then used a score threshold of 2.5 to select cells in the cluster of interest. Differential expression analysis was performed between the cell cluster of interest and all other remaining cells using the rank_genes_groups function in scanpy (Wilcoxon test). Details of the analysis can be found in our jupyter notebook B_cell_scRNAseq.ipynb. To reconstruct CDR3 regions of single cells, we demultiplexed bulk fastq files into single cell fastq files with *scruff* package in R ^72^, mapped each fastq to GRCm38.p6 genome and obtained gene counts with STAR v2.7.2b ^53^, and reconstructed CDR3 regions of Ig kappa chain from individual cells using mixcr ^73^. Clone sizes were calculated as percent of templates supporting the current Ig kappa chain to the total number of reconstructed templates of Ig kappa chain for the sample. Cells with fewer than 200 gene counts were removed. Raw gene counts were log-transformed and normalized using the *edgeR* package in R ^68^. Clonality signature score was calculated for each cell as transcript level of top 50 genes minus transcript level of bottom 50 genes ranked by p-value and taken from our bulk RNA-seq regressed against clone sizes (Supplementary Table 3). tSNEs were calculated using the *M3C* package in R ^74^.

To identify ABC and ACBC clusters in human donors we translated the generated signatures for ABC and ACBC clusters from mouse to human genes by mapping the Ensembl IDs. Single-cell RNA sequencing data was acquired from a recent study available on cellxgene (https://cellxgene.cziscience.com/collections/0a839c4b-10d0-4d64-9272-684c49a2c8ba) and the corresponding metadata from the Gene Expression Omnibus (GEO) database (https://www.ncbi.nlm.nih.gov/geo/query/acc.cgi?acc=GSE158055). Cells with available BCR sequences ("BCR single cell sequencing" == "Yes") were selected from 19 healthy donors and retained only B cells or cells with an assigned BCR clone ID.

Data preprocessing was performed using scanpy ^71^. Genes were filtered using min_cells=50, and cells were filtered using min_genes=200. Then a scVI ^75^ model was trained with default parameters (n_hidden=128, n_latent=10, n_layers=1) for 50 epochs using 6,000 highly variable genes calculated with sc.pp.highly_variable_genes (seurat_v3 flavor and batch_key=’Sample type’) and the marker genes from the ABC and ACBC signatures. Neighbors were generated with sc.pp.neighbors using the scVI latent space, and UMAP was calculated with sc.tl.umap. In the resulting density plot, calculated with scanpy’s ^71^ sc.pl.embedding_density, clonal cells were marked as ’yes’ when clones had >1 counts, and clone frequency was directly obtained from the metadata. ’abc’, ’acbc’, and ’myc_targets’ scores were generated using the ’aucell’ module from pyscenic ^76^. scVI normalized expression matrix was used and generated with vae.get_normalized_expression(library_size=10e4, n_samples=1) for expression calculations. For ’abc’ and ’acbc’, scores were independently calculated for upregulated and downregulated genes from the mouse signature and then subtracted the downregulated score from the upregulated score. ’myc_targets’ were downloaded from the TRRUST v2 database (https://www.grnpedia.org/trrust/) and filtered for genes that are activated by c-Myc. The values were plotted using sc.pl.umap with vmin=0.0, vmax=’p95’, and color_map=’inferno’. For the Ig chain plots, ’acbc’ cells were defined as cells with ’is_clonal’, ’acbc’ > 0.2, and ’myc_targets’ > 0.3. Subsequently, Ig chains were plotted derived from the metadata.

### Reconstruction of Ig CDR3 regions

Genomic CDR3 regions of Ig heavy chains were analyzed with Immunoseq (Adaptive Biotechnologies). CDR3 regions were reconstructed from RNA sequencing raw data using mixcr software ^77^ with recommended settings for transcriptome data. Filtering of reconstructed regions and diversity analysis was done with VDJtools software ^78^.

### Immunofluorescence microscopy

B cells were FACS-sorted at 500 thousand cells per well and incubated with poly l-lysine treated coverslips for 1 hour in 24 well plates. Cells were permeabilized with 0.1% Triton X-100 2 times for 30 seconds, fixed in 3.7% PFA in PBS for 10 minutes and washed three times with PBS, incubated with the blocking buffer until further analysis (1% BSA, 0.1% Triton X-100 in PBS). Samples were incubated with primary antibodies overnight at 4 °C (1:100, Abcam #ab32072), then washed with PBS five times and incubated overnight with Alexa Fluor 568 - conjugated anti-rabbit secondary antibodies (1:500, Biotium cat#20098) and DAPI dye. Cells were washed with PBS and mounted with ProLong Diamond antifade (Thermo Fisher Scientific). Samples were imaged using Leica SP8 confocal microscope. Images were analyzed with ImageJ ^79^. Raw Z-stacks were converted to the maximum intensity projection images. Nuclei and cell borders were detected using manual thresholding and “Analyze Particle” function. All crowded groups and not-round shaped cells were manually removed from the analysis.

### Reduced representation bisulfite sequencing (RRBS)

Libraries were prepared and sequenced as in ^80^. Bisulfite sequence reads were trimmed by TrimGalore v0.4.1 and mapped to the mouse genome sequence (mm10/GRCm38.p6) with Bismark v0.15.0 ^81^. We kept CpG sites that were covered by five reads or more. Promoter regions were determined as the [-1500, +500] bp from the transcription start site (following the direction of the transcription) taken from Ensembl annotation file (Mus_musculus.GRCm38.100.chr.gtf). The start and end positions of gene bodies were taken from the Ensembl gene predictions (Mus_musculus.GRCm38.cds.all.fa). The mean methylation levels were calculated for regions that have at least 5 covered CpG sites with average methylation level above 1%. To determine ribosomal DNA methylation (rDNAm) age, we developed a blood rDNAm clock in a similar way as described in ^82^. Briefly, we applied ElasticNet regression on ribosomal DNA (BK000964.3) CpG methylation levels of 153 control fed C57BL/6 blood samples with an age range from 0.67 to 35 months (GSE80672). We identified the solo-WCGWs (i.e. CpGs flanked by an A or T on both sides) in partially methylated domains (PMDs) based on available coordinates for the mouse mm10 genome (https://zwdzwd.github.io/pmd) ^33^.

### Prediction of biological transcriptomic age

To assess biological transcriptomic age (tAge) of B cells from young mice, old mice and mice with lymphoma, we applied a mouse multi-tissue clock of lifespan-adjusted age developed based on gene expression signatures of aging identified previously ^83^. For RNAseq data, we filtered out genes with low number of reads, keeping only the genes with at least 10 reads in at least 50% of the samples, which resulted in 10,670 detected genes according to Entrez annotation. Filtered data was then passed to log transformation and scaling. The missing values corresponding to clock genes not detected in the data were omitted with the precalculated average values. Resulted tAge values were centered based on median tAge of B cells from young animals. Pairwise differences between average tAges across groups were assessed using independent samples two-tailed t-test.

### Longitudinal blood collection and analysis

Mice were anesthetized with isoflurane and then locally with topical anesthetic, restrained, and approximately 100 ul of blood was collected from mouse tails into EDTA-coated tubes (BD). Blood was incubated on ice until further analysis (2-3 hours), then mixed with 1 ml of red blood cell lysis buffer and centrifuged at 250g for 10 minutes at 4°C. Pellets were washed once with a FACS buffer (PBS with 1% FBS), split equally into 2 tubes and incubated for 20 minutes at 4°C with antibodies against B cell, T cell and myeloid cell markers, or follicular, marginal zone, and plasma cell markers. Stained cells were washed again, resuspended in 200 ul of FACS buffer and analyzed with FACS, with 20,000 events being recorded. Dead cells were gated by DAPI staining. Cell size was measured with forward scatter.

### Longitudinal blood scores

We defined the FSC score as the difference between the last measurement of mean B cell size prior to death (if mouse died) or B cell size at the given round (if mouse was alive) and the mean B cell size of young mice at the same round. CD21.score was calculated as delta(CD21-CD23-) - delta(Follicular). Delta(CD21-CD23-) was calculated as percentage of CD21^-^CD23^-^ B cells of total B cells before death (if mouse died) or percentage of CD21^-^CD23^-^ B cells at the given round (if mouse was alive) minus percentage of CD21^-^CD23^-^ B cells in young mice at the same round. delta(Follicular) was calculated the same way for the percentage of follicular B cells of B cells. The myeloid score was calculated as delta(Myeloid) - delta(B cells). delta(Myeloid) was calculated as percentage of myeloid cells of CD45^+^ cells before death (if mouse died) or percentage at the given round (if mouse was alive) minus the percentage of myeloid cells in young mice at the same round. delta(B cells) was calculated the same way for the percentage of B cells of CD45+ cells. A higher CD21.score indicates a higher proportion of CD21^-^CD23^-^ B cells to total B cells and/or lower proportion of follicular B cells to total B cells. A higher myeloid score indicates a higher proportion of myeloid cells to CD45+ cells and/or lower proportion of B cells to CD45+. A higher FSC score indicates a greater increase of B-cell size.

### Cell culture

Freshly FACS-sorted cells were plated into 96 wells at 400,000 cells per well for each cell type and cultured for 24-48 hours in 200 ul of RPMI medium with 10% FCS (Gibco), 2 mM glutamine (ThermoFisher), 1% oxaloacetic acid (15 mg/ml), 5 mg/ml sodium pyruvate (ThermoFisher), 1% non-essential amino acids (ThermoFisher), and 50 μM 2-ME (Sigma). Where follicular B cells were incubated alone, 800,000 cells were plated to maintain the same cell density between control and treatment wells. In the case of CD22 co-incubation, follicular B cells or ABC from the spleens of 25-month-old mice were freshly FACS-sorted and plated at 100,000 cells per well density into 96-well plates in the same media described above. InVivoMAb anti-mouse CD22 antibody (Bio X Cell, cat# BE0011) was added for the final concentration of 5 μg/ml. Cells were cultured overnight and analyzed with flow cytometry. After co-incubation, cells were centrifuged at 250 g for 10 minutes at 4°C with slow deceleration. The media were carefully removed leaving ∼50 ul, cells were washed once in 100 ul of FACS buffer, then resuspended in 100 ul of AB solution (1:100) and incubated at 4°C for 20 minutes protected from light, washed again and resuspended in 200 ul of FACS buffer, and filtered through 40 µm Falcon Cell Strainers (Corning). Twenty to one hundred thousand cells were recorded, and follicular B cells were gated as CD19^+^CD21^int^CD23^+^ and FACS-sorted into 300 ul of Trizol or analyzed with flow cytometry analyzer. RNA was extracted with Direct-zol RNA Microprep (Zymo Research).

### Necropsy analysis

Mice were euthanized with CO_2_ followed by cervical dislocation. The chest and abdomen were opened, and the body was immersed into formalin solution and stored at 4°C until further analysis. For necropsy analysis all organs, including small endocrine organs, were dissected, trimmed at 5 mm thickness and embedded in paraffin blocks. Paraffin blocks were sectioned at 5 μm and stained with hematoxylin and eosin. The slides were examined blindly by a pathologist. Lymphoma was diagnosed when multiple solid tissues contained large uniform sheets of atypical lymphocytes with large nuclei. Lymphocytic hyperplasia was diagnosed when any solid tissue had small infiltrates of atypical lymphocytes with large nuclei.

### Pseudo-time analysis

Pseudo-time analysis is a computational technique used in single-cell transcriptomics to infer the developmental trajectory of cells from gene expression data. Here the pseudo-time analysis was performed using *slingshot* R package ^84^. The FO/MZ B cell cluster was assigned to be the initial cluster and the pseudo-time inference was performed using the UMAP embedding and default parameters. The result is visualized using custom R script and ggplot2.

### Data analysis and availability

All data were analyzed and plotted with R in Rstudio. RNA sequencing, DNA methylation and proteomics data were preprocessed and analyzed for differential changes and GSEA with *limma* ^85^, *edgeR* ^68^ and *clusterprofiler* ^86^ packages. All p-values for group means comparisons were calculated with two-tailed Student t-test, unless otherwise specified. Correlations between two variables were evaluated using Pearson’s correlation coefficient. PCA analysis was done with the factoextra package. Color schemes are from the ggsci package. Raw reads for RNA-sequencing, whole exome sequencing, and RRBS are available at SRA (PRJNA694093).

## Acknowledgments

The authors thank members of the Gladyshev laboratory for discussion. Supported by Max Kade Foundation to JPC and by NIH grants to VNG.

## Author Contributions

JPC and AVS conceived the project, designed and performed experiments, analyzed the data and drafted the manuscript. AB, OSS, JAP, CK, APP, MM, MM, and YH performed experiments, analyzed the data and revised the manuscript. GL, SPG, and JMS provided research materials, assisted with experimental design and revised the manuscript. JPM and VNG interpreted the data, designed experiments, provided research materials, and revised the manuscript. VNG obtained funding and supervised the overall project.

