## Supplementary material for "Integrative analyses uncover mechanisms by which aging drives B cell lymphoma": Castro and Shindyapina et al 2023 Supp material 2023.pdf

#### Figure S1

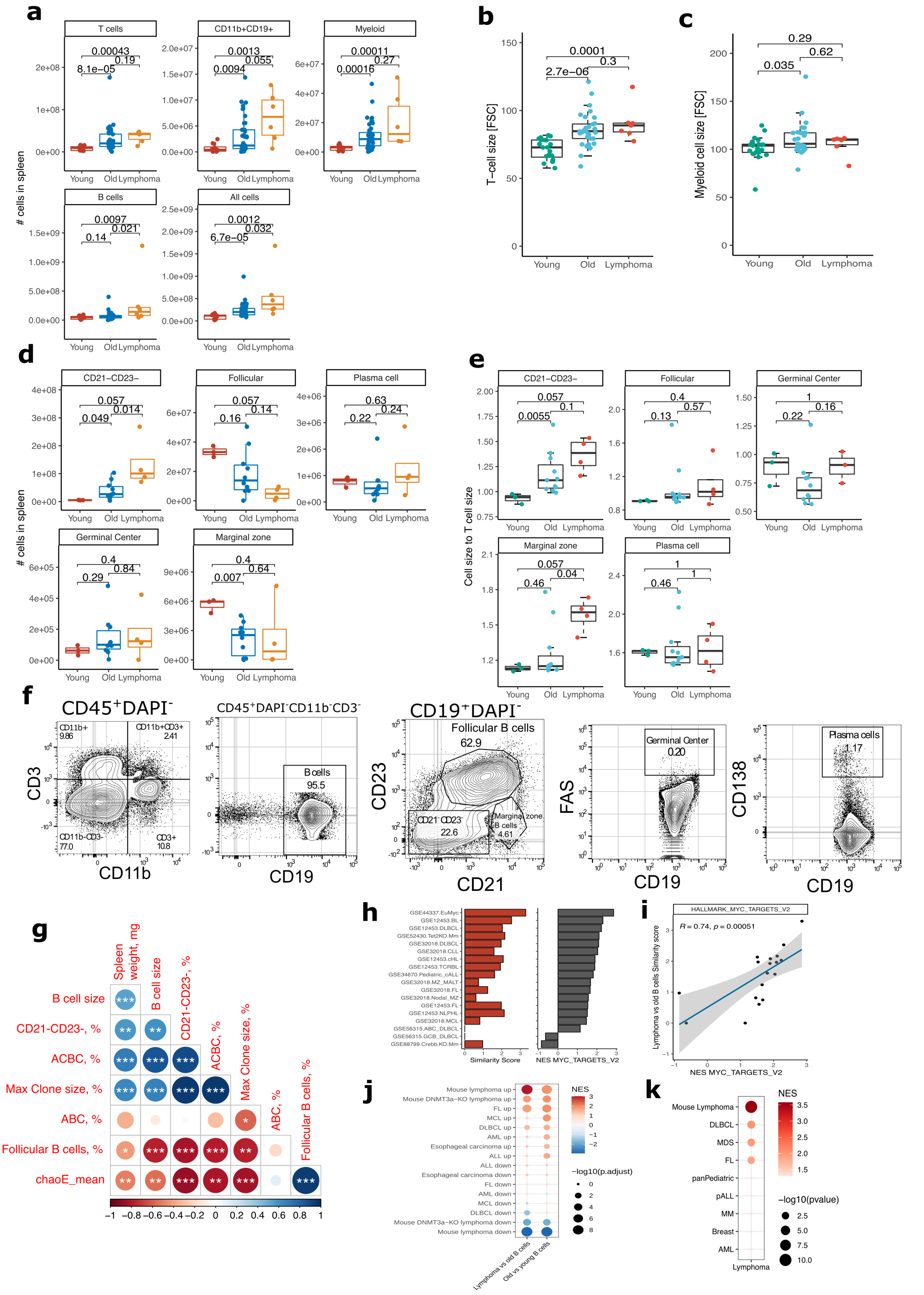

**Supplementary Figure 1. Age-related changes in cell size and immune cell composition in C57BL/6 mice.** (a) Number of immune cells in the spleens of young, old and old lymphoma-bearing mice as measured by FACS and calculated based on the total number of cells in the spleen. (b) Mean size of T cells or (c)

myeloid cells in the spleens as measured by FACS forward scatter. (d) Number of B cells in the spleens of young, old and old lymphoma-bearing mice as measured by FACS and calculated based on the total number of cells in the spleen. (e) Mean cell size in the spleens of young, old and old lymphoma-bearing mice as measured by FACS normalized to T cell size from the same donors. (f) Gating strategy for B cells. (g) Gating strategy for follicular, marginal zone, CD21-CD23-, germinal center B cells and plasma cells. (g) Correlation between age-related phenotypes of B cells measured in old C57BL/6 mice. Scale bar is a Pearson correlation coefficient. Asterisks indicate statistical significance. **** p-value < 0.0001, *** p-value < 0.001, ** p-value < 0.01, * p-value < 0.05. (h) Similarity score of the transcriptomic signature of mouse cancerous B cells versus publicly available datasets of different human and mouse cancers (left plot). Enrichment scores for the Hallmark MYC targets pathway for human and mouse cancers (right plot). (i) Correlation between the similarity score from (h) on the left plot and enrichment score (NES) from (h) on the right plot. R is a Pearson correlation coefficient. (j) Enrichment of differentially methylated promoters found in B cells from old or lymphoma-bearing mice against those found in human cancers. ‘Mouse lymphoma up’ and ‘Mouse lymphoma down’ gene lists were created from our DNA methylation dataset and used as positive control in the analysis. (k) Enrichment of genes mutated in mouse lymphoma against these mutated in human cancers. ‘Mouse lymphoma’ gene list was created from our genome sequencing dataset and used as a positive control in the analysis

### Figure S2

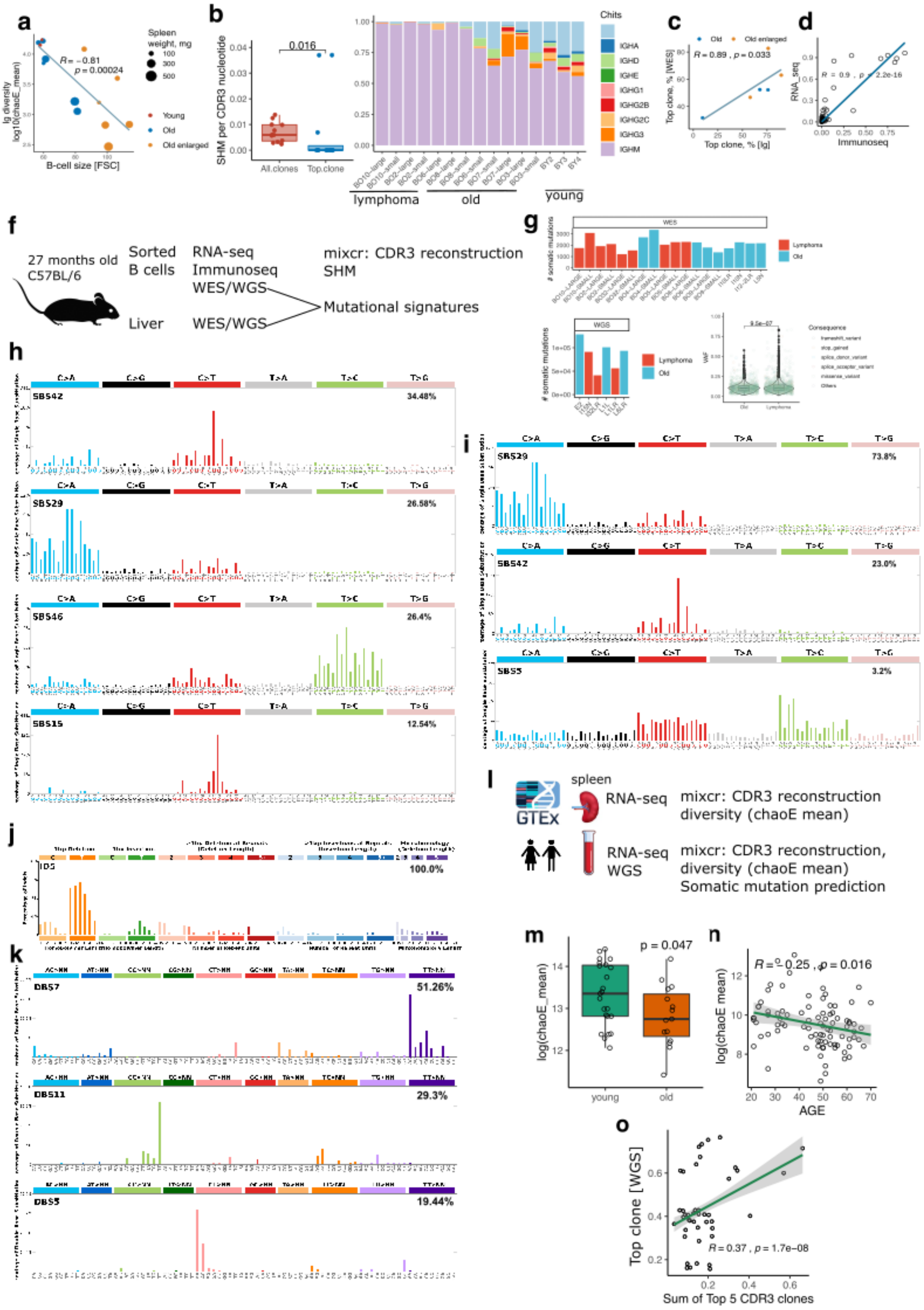
a
b
c
d
f
g
h
i
j
k
l
m
n
o
Spleen weight, mg
Young
Old
Old enlarged
B-cell size [FSC]
lg diversity log10(chaoE_mean)
R = -0.81
p = 0.00024
SHM per CDR3 nucleotide
0.016
All.clones
Top.clone
Chits
IGHA
IGHD
IGHE
IGHG1
IGHG2B
IGHG2C
IGHG3
IGHM
lymphoma
old
young
R = 0.89, p = 0.033
Top clone, % [WES]
Top clone, % [Ig]
R = 0.9, p < 2.2e-16
RNA_seq
Immunoseq
27 months old
C57BL/6
Sorted
B cells
RNA-seq
Immunoseq
WES/WGS
Liver
WES/WGS
mixcr: CDR3 reconstruction
SHM
Mutational signatures
WES
WGS
Lymphoma
Old
### somatic mutations
SBS42
SBS29
SBS46
SBS15
SBS5
34.48%
26.58%
26.4%
12.54%
73.8%
23.0%
3.2%
ID5
100.0%
DBS7
DBS11
DBS5
51.26%
29.3%
19.44%
GTEx
spleen
RNA-seq
mixcr: CDR3 reconstruction
diversity (chaoE mean)
RNA-seq
WGS
mixcr: CDR3 reconstruction,
diversity (chaoE mean)
Somatic mutation prediction
p = 0.047
log(chaoE_mean)
young
old
R = -0.25, p = 0.016
log(chaoE mean)
AGE
Top clone [WGS]
R = 0.37, p = 1.7e-08
Sum of Top 5 CDR3 clones

**Supplementary Figure 2. Mouse and human B cells clonally expand with age, which leads to reduced Ig diversity.** (a) Correlation between B-cell size as measured by FACS and the size of top B-cell clone as measured based on reconstruction of CDR3 regions from RNA-seq data to estimate Ig (ChaoE_Mean) from FACS-sorted B cells from young and old mice. (b) (left) Proportion of somatic mutations per nucleotide within the CDR3 region in all clones compared to largest clones. (right) IgH isotype distribution across young, old and lymphoma samples. Light blue means “not detected”. (c) Correlation between top clone size (Ig) estimated from CDR3 sequencing and top clone size estimated by VAF from whole exome sequences. (d) Correlation of clone size for the same clones reconstructed from RNA and from genomic sequencing of CDR3 regions. (e) Scheme of the analysis of sorted splenic B cells from 27 months old mice by RNA-seq, Immunoseq and WES/WGS. Liver WES/WGS were used as a control to exclude germline somatic mutations. (f) Number of somatic mutations per sample identified with whole exome (upper plot) or genome (bottom, left plot) sequencing. (g) (h) (i) and (j) reconstructed mutational signatures of total somatic mutations in B cells of old mice. (k) Scheme of the analysis of the GTEx dataset. RNA sequences from spleen and blood were used to reconstruct CDR3 regions, estimate Ig diversity (chaoE mean), and WGS of blood from the same subjects was used to predict somatic mutations. (m) Ig diversity (chaoE_mean) in spleens of young and old GTEx subjects calculated from RNA-seq data. P-value was calculated with a two-sided Student t-test. (n) Correlation between Ig diversity (chaoE_mean) in blood of GTEx subjects and their age, in years. (o) Correlation between top clone (WES) and Ig clone sizes (sum of top 5 CDR3 clones) in blood of GTEx subjects.

#### Figure S3

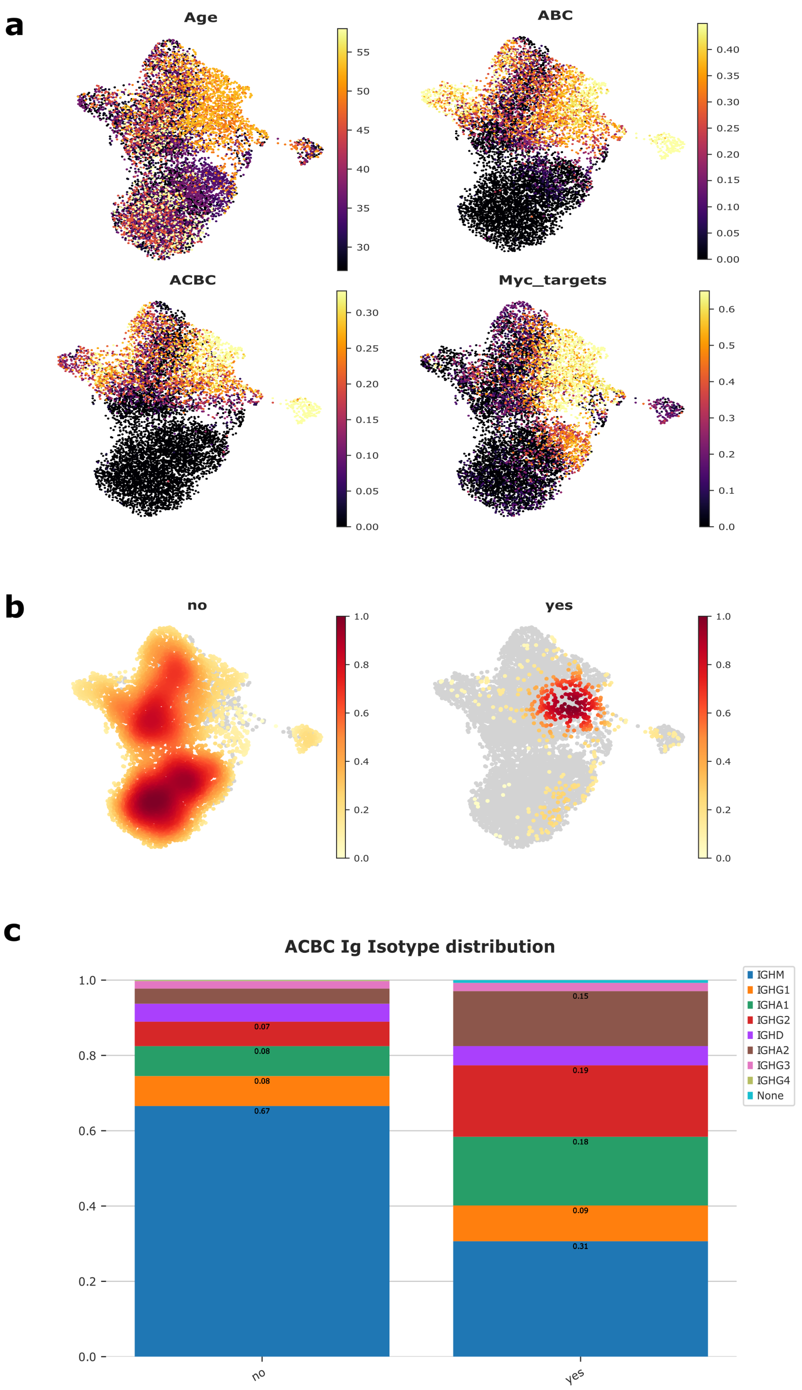


**Supplementary Figure 3. Increased B cell clonality and enrichment for ABC and ACBC signatures in aged human donors.** (a) UMAPs showing ABC, ACBC and Myc targets enrichment across age gradient. (b) UMAPs showing the distribution of cells with “no” and “yes” for B cell clonality (c) IgH isotype distribution in ACBC enriched B cells. Single-cell RNA sequencing data was acquired from [39] and the corresponding metadata from the Gene Expression Omnibus (GEO) database (GSE158055). Cells with available BCR sequences ("BCR single cell sequencing" == "Yes") were selected from 19 healthy donors and retained only B cells or cells with an assigned BCR clone ID.

#### Figure S4

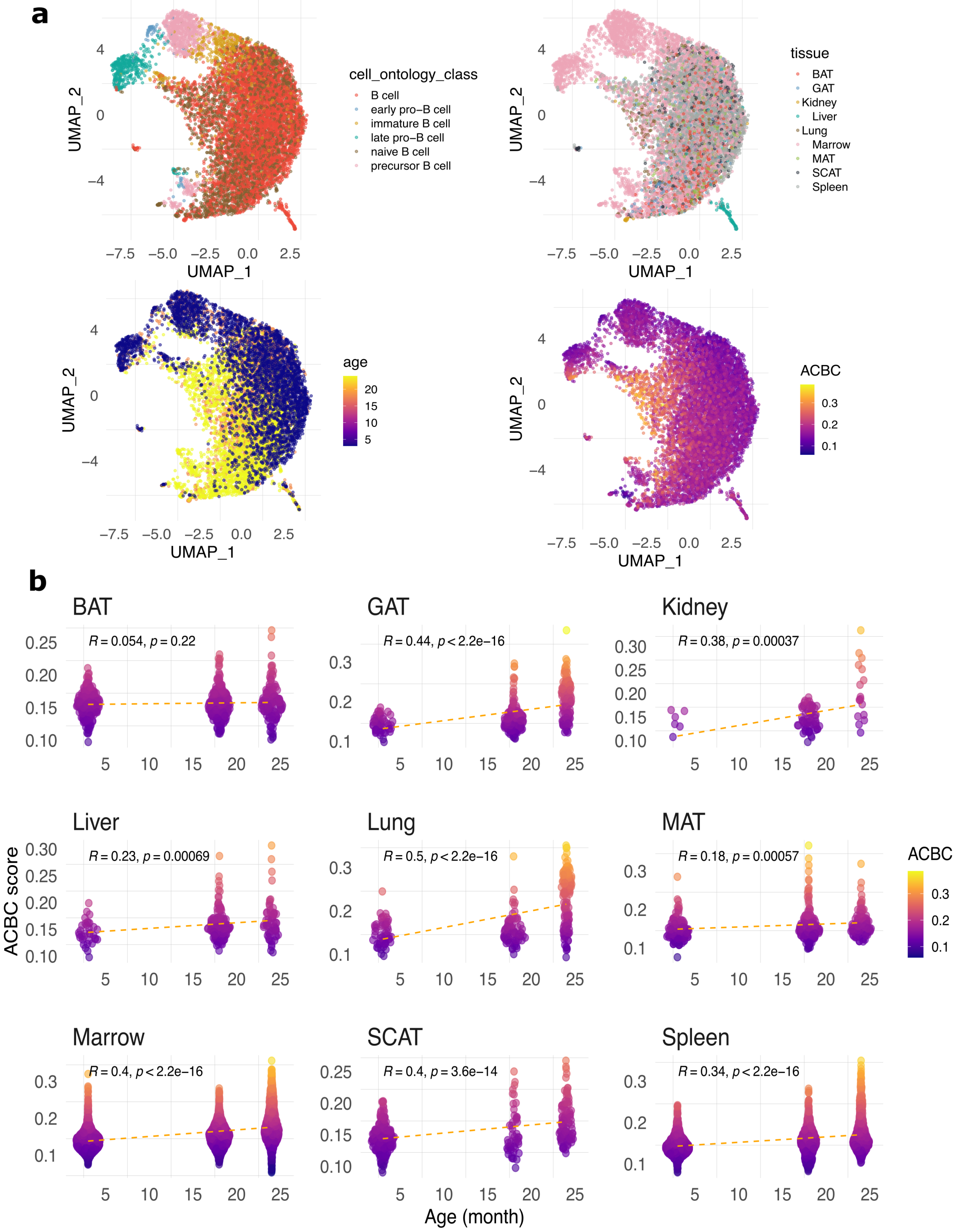


**Supplementary Figure 4. Age-associated clonal B cells infiltrate peripheral tissues.** (a) UMAP plots of single-cell RNA-seq from many combined tissues across mouse lifespan in months. Data was obtained from *Tabula Muris Senis* dataset [37] (b) ACBC signature enrichment in B-cell populations in a multitude of tissues such as Brown adipose tissue (BAT), Gonadal Adipose Tissue (GAT), Kidney, Liver, Lung, Mesenteric adipose tissue (MAT), Bone marrow, Subcutaneous adipose tissue (SCAT) and spleen. All data was extracted from *Tabula Muris Senis* dataset[40]. Cell signature scores were performed by UCell and statistical analysis was performed based on Mann Whitney U statistic [87].

#### Figure S5

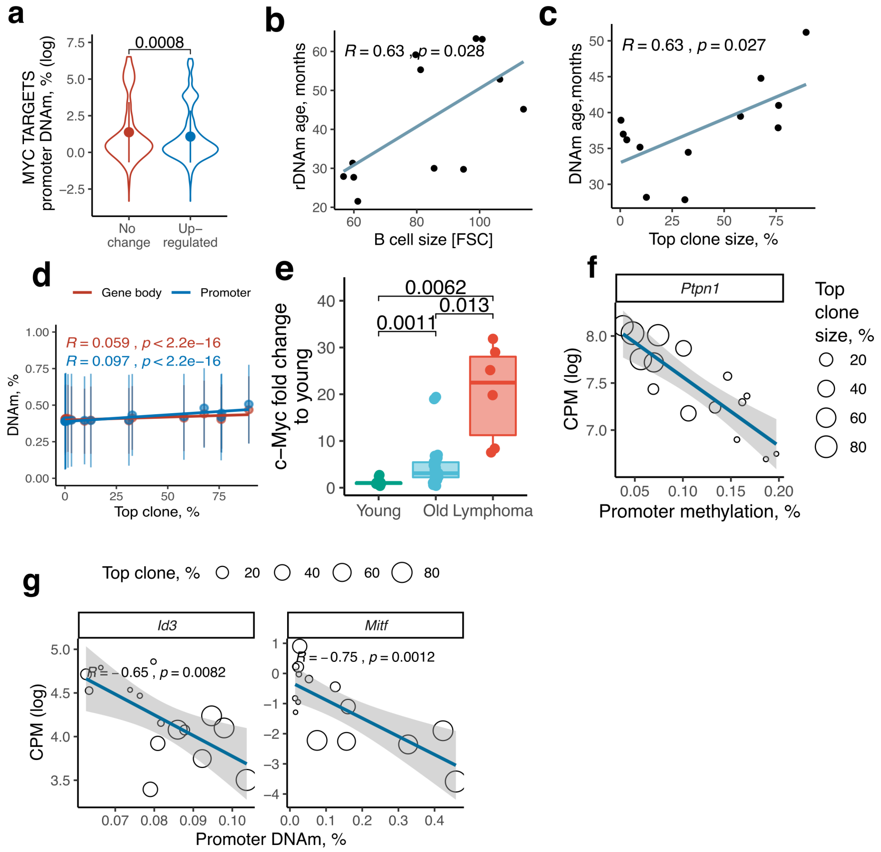


**Supplementary Figure 5. Clonal expansions of B cells are supported by c-Myc and DNA methylation alterations.** (a) Mean DNA promoter methylation levels for genes encoding c-Myc targets. (b) Correlation between mean B cell size and the rDNA methylation age. (c) Correlation between the DNA methylation age calculated by the blood epigenetic clock[36] and the size of top B cell clones. (d) Correlation of top Ig clone size and mean global DNA methylation of promoters and gene bodies in B cells of old mice. (e) c-Myc fold change in young, old and lymphoma FACS-sorted B cells normalized for DNA concentration and measured by qRT-PCR, p-values were calculated with a two-sided Student t-test (f) Correlation between *Ptpn1* gene expression and mean DNA methylation of its promoter, dots are sized by size of top clone of B cells in a sample. Correlations between two variables were evaluated using Pearson's correlation coefficients (g) Same as (f) but for *Id3* and *Mitf* genes.

#### Figure S6

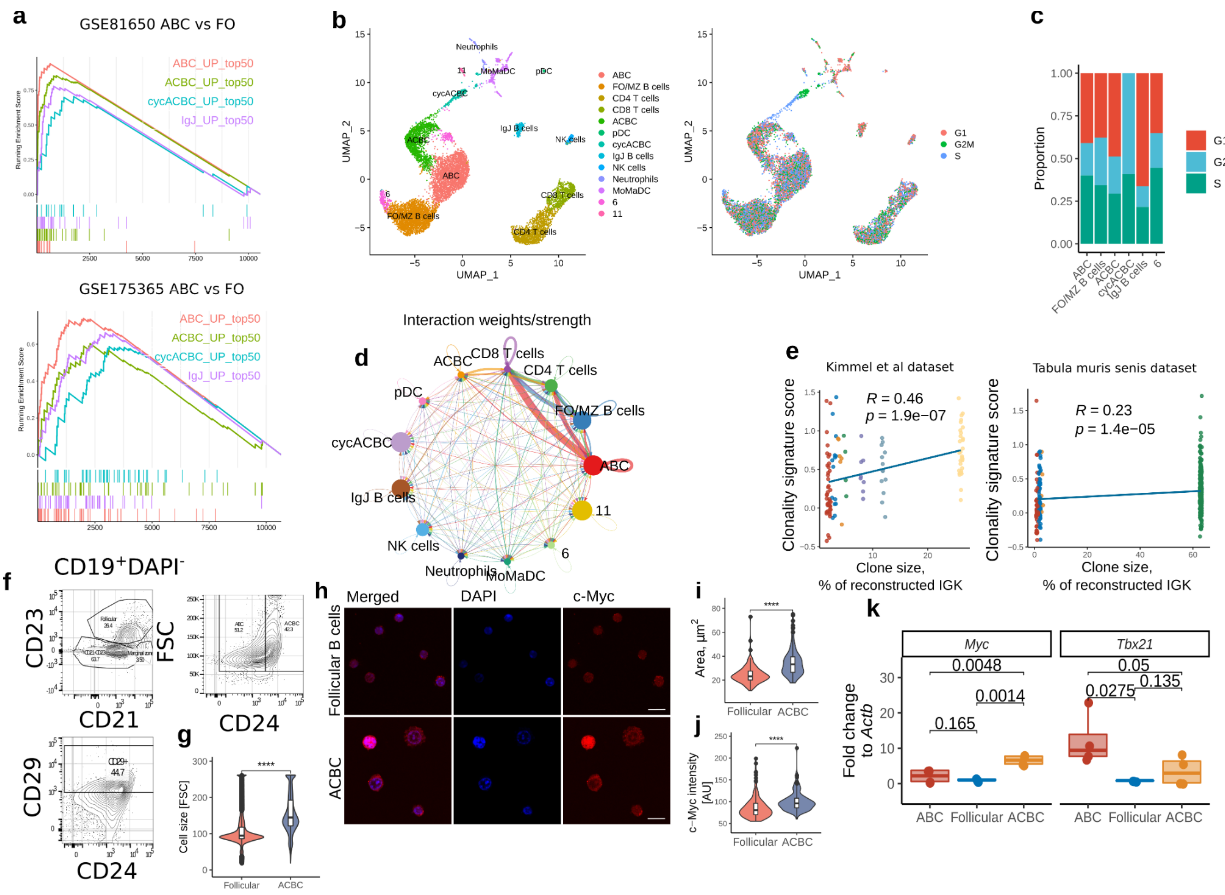


**Supplementary Figure 6. Single-cell RNA sequencing uncovers a clonal B-cell population and its relation to age-associated B cells (ABC).** (a) Enrichment plots of gene expression changes in ABC based on bulk public RNA-seq datasets for signatures from age-related B cell populations based on single-cell RNA-seq. (b) UMAP plots of single-cell RNA-seq of the spleens of old mice from the combined Kimmel dataset[36] and *Tabula Muris Senis* dataset[37] (left plot). Immune cell enrichment in mitotic genes clustered according to G1, G2M and S phases (right plot). (c) Proportion of cells predicted to be in G1, G2M or S mitotic phases across B cell populations. (d) Predicted interaction of immune cells from single-cell RNA-seq using CellChat. (e) Correlation between clonality signature score estimated for each cell and clone size (% of reconstructed IgK) to which it belongs using two publicly available datasets (39, 40). (f) Gating strategy for ABC and ACBC populations. (g) Cell sizes measured by FSC of follicular and ACBC in the spleens of young (n=4) and old (n=10) mice. (h) Representative images of follicular and ACBC cells immunolabeled for *c-Myc* and for DNA with DAPI. Scale bar, 10 µm. (i) Quantification of cell sizes and (j) mean *c-Myc* intensity for follicular and ACBC estimated from confocal images for two independent experiments (k) *c-Myc* and *Tbx21* fold change to *Actnb* measured with qRT-PCR in splenic follicular B cells, ABC and ACBC FACS-sorted from 25-27-month-old mice, p-value was calculated with a one-sided Student t-test.

#### Figure S7

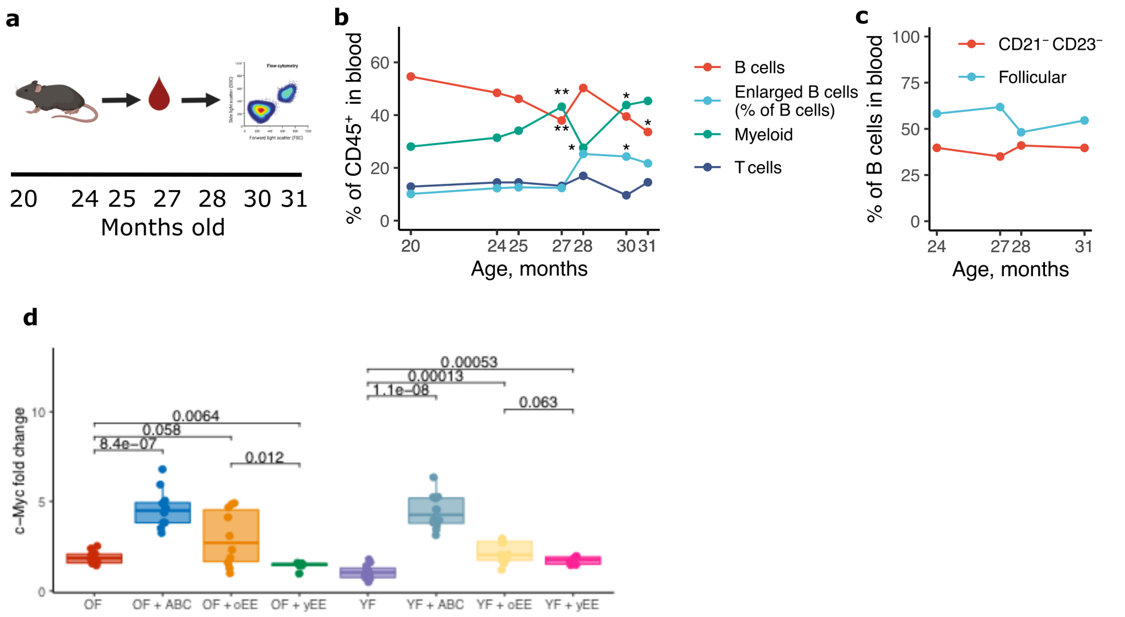


**Supplementary Figure 7. Longitudinal analysis reveals B-cell size and CD21$^-$CD23$^-$ B cells as predictors of lifespan, and co-culture experiments point to their origin.** (a) Schematic representation of the longitudinal experiment. Blood was collected from the tails of C57BL/6 female mice across seven timepoints for FACS analysis. (b) and (c) changes in blood cell composition with age in 25 old mice. (d) *c-Myc* transcript levels measured by RT-PCR in young (YF) and old follicular (OF) B cells incubated with i) age-associated B cells (ABC) from old spleens, ii) immune cells (CD45$^+$) from young spleens excluding B cells (yEE), or iii) immune cells (CD45$^+$) from old spleens excluding B cells (oEE). Results shown are from three independent experiments, 3-4 biological replicas per experiment.

#### Figure S8

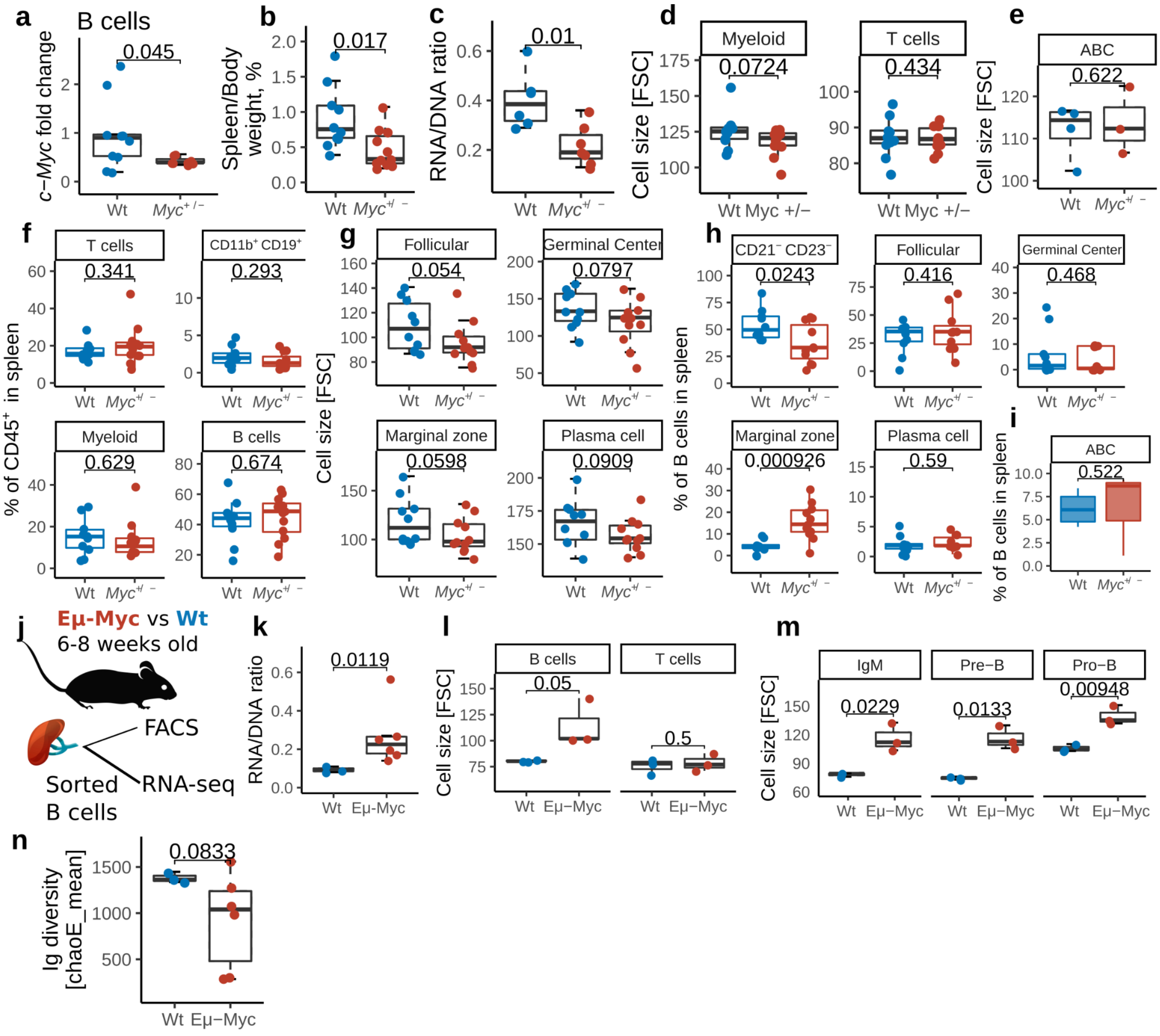


**Supplementary Figure 8. Effects of *c-Myc* deficiency and overexpression on immune cell composition and premalignant phenotypes of B cells.** (a) c-Myc expression fold change normalized to DNA concentration in B cells, (b) spleen weight normalized by body weight, and (c) the RNA to DNA concentration ratio in B cells, (d-f) splenic cell sizes from *Myc*$^{+/-}$ mice and their wild type (Wt) siblings. (g) Cell composition of immune cells (CD45$^{+}$) and B-cell populations (h,i) in the spleens of old Wt and *Myc*$^{+/-}$ mice, gated as in Fig. S1f-g. (j) Analysis of Emu-*Myc* mice. (k) Total RNA/DNA ratio in FACS-sorted splenic B cells from non-carrier (NC) controls and Emu-*Myc* mice. (l) Mean size of B and T cells measured by FACS in the spleens of NC and Emu-*Myc* mice. (m) Mean cell size of B-cell populations measured by FACS in the bone marrow of NC and Emu-*Myc* mice. (n) Ig diversity (chaoE_mean) of splenic B cells from NC and Emu-*Myc* mice, calculated from RNA-seq data using mixcr and VDJtools software.
